# Single Cell Epigenetics Reveal Cell-Cell Communication Networks in Normal and Abnormal Cardiac Morphogenesis

**DOI:** 10.1101/2022.07.25.501458

**Authors:** Sanjeev S. Ranade, Sean Whalen, Ivana Zlatanova, Tomohiro Nishino, Benjamin van Soldt, Lin Ye, Angelo Pelonero, Langley Grace Wallace, Yu Huang, Michael Alexanian, Arun Padmanabhan, Barbara Gonzalez-Teran, Pawel Przytycki, Mauro W. Costa, Casey A. Gifford, Brian L. Black, Katherine S. Pollard, Deepak Srivastava

## Abstract

Communication between myriad cell types during organ formation underlies proper morphogenesis^1^. In cardiac development, reciprocal signaling between mesoderm progenitors and neural crest cells is essential, and its disruption leads to congenital heart malformations, the most common human birth defect. However, mechanistic interrogation of temporal gene networks and *cis* regulatory elements in this crosstalk is limited^2,3^. Here, we integrated single cell chromatin accessibility and transcriptomics to establish an unbiased and temporal epigenomic map of the embryonic mouse heart over multiple stages and developed machine learning models to predict enhancers for heart and neural crest. We leveraged these advances to determine the consequences of dysregulated signaling at single cell resolution caused by deletion of TBX1, a transcription factor that causes morphogenetic defects of the cardiac outflow tract in humans and functions non-cell autonomously in cardiac mesodermal progenitors to direct pharyngeal neural crest differentiation^4–6^. Loss of Tbx1 in mice led to broad closure of chromatin regions enriched in cardiac progenitor transcription factor motifs within a narrow subset of cardiac mesodermal progenitors and correlated with diminished expression of numerous members of the fibroblast growth factor, retinoic acid, Notch and Semaphorin pathways. In affected progenitors, ectopic accessibility and expression of posterior heart field factors in the anterior heart field suggested impaired axial patterning. In response, a subset of cardiac neural crest cells displayed epigenomic and transcriptional defects, indicating a failure of differentiation corresponding to dysregulation of the anterior-posterior gradient of pharyngeal Hox gene expression. This study demonstrates that single-cell genomics and machine learning can generate a mechanistic model for how disruptions in cell communication selectively affect spatiotemporally dynamic regulatory networks in cardiogenesis.

---

Birth defects occur in ~6% of live births and are the result of abnormalities in morphogenesis^7^. Congenital heart defects (CHD) are the most common form, present in ~1% of newborns, with more severe deformities responsible for ~10% of spontaneous first trimester fetal loss^2,8^. Disruption of reciprocal signaling between cells that form the cardiac outflow tract, which facilitates blood flow out of the heart, underlies ~35% of CHD^8^. Essential signaling and transcription factors involved in cardiac lineage specification are well-defined, and single cell RNA-sequencing (scRNA-seq) studies have begun to reveal the heterogeneity of cell types based on unique transcriptomes^9^. However, annotation of *cis* regulatory elements controlling gene expression in cardiac development has been achieved primarily through *in vitro* differentiation systems, and the underlying epigenomic steps that distinguish discrete cell types and dictate morphogenetic cues through cell-cell communication *in vivo* are poorly understood^3,10–12^. Mapping such cell type-specific *cis* regulatory elements will ultimately be necessary to interpret non-coding genetic variation that may contribute to disease.

During cardiogenesis, two distinct pools of mesodermal progenitor cells, termed first and second heart fields (SHF), predominantly contribute to the heart chambers and valves, while neural crest-derived cells shape vessels of the outflow tract. Multipotent neural crest cells that emigrate from the neural tube receive signals from SHF progenitors in the pharyngeal arches and differentiate along an axis such that cells in the rostral pharyngeal arches (PA1 and PA2) give rise to craniofacial derivates, whereas cells within the caudal arches (PA3-6) affect cardiac patterning^13^. Ablation of cardiac neural crest cells causes outflow tract anomalies, establishing the necessity of this lineage for heart formation^14^. In humans, 22q11 deletion syndrome, the most common genetic deletion condition involving a ~3 Mb region of chromosome 22, results in craniofacial and cardiac outflow tract abnormalities and is considered a neurocristopathy^5^. TBX1, a transcription factor located within the deleted region and a main driver of the cardiac and craniofacial defects, is expressed in multiple cell types within the heart and pharyngeal arches, but not within neural crest-derived cells, suggesting a non-cell autonomous function^5,15^. Although previous studies have implicated mis-direction of neural crest migration, the precise cell types and dysregulated gene networks in relevant cells underlying disrupted neural crest differentiation due to loss of Tbx1-dependent signaling are not known^15–17^.

Recent advances in single cell assay for transposase accessible chromatin with sequencing (scATAC-seq) and integration with scRNA-seq have transformed our ability to capture and correlate epigenomic and transcriptomic states of single cells in embryonic development, allowing dissection of non-cell autonomous functions of gene networks^18,19^. Loss of function mouse models of Tbx1 recapitulate most of the cardiac and craniofacial developmental defects observed in humans and serve as a model for mechanistically interrogating how disruption of cell-cell communications affect morphogenesis^20^. In this study, we undertook an integrated multimodal single cell approach to map the *cis* regulatory elements (cREs) of all cells during normal early heart development, developed novel machine learning models for enhancer prediction from scATAC-seq data, and then showed how loss of Tbx1 disrupts regulatory networks within subsets of both cardiac and neural crest cells leading to axial patterning abnormalities and cardiac morphogenetic defects.

## Results

### Chromatin Accessibility Map of Mouse Heart Development at Single Cell Resolution

To define regulatory elements and transcriptomes during cardiac development at single cell resolution, we employed scATAC-seq together with scRNA-seq^9,18,19^. Micro-dissected tissue from wild type C57Bl/6J mouse heart and pharyngeal arch regions were captured from five stages of cardiogenesis between E7.75 – E11.5, with earlier stages (E7.75 – E9.5) containing all regions and later stages (E10.5 and E11.5) prioritizing regions dorsal to the heart enriched for SHF progenitors and neural crest cells (Table S1). For scATAC-seq, we used ArchR to process all samples and identified 41 epigenetic clusters containing 64,956 cells; for scRNA-seq, we used Seurat to process and group 61,530 cells into 46 transcriptomic clusters (Extended Data Fig. 1a)^21,22^. Cell type-enriched marker genes from scRNA-seq were then assessed in scATAC-seq using gene scores, which are imputed chromatin accessibility values within and around the gene body, facilitating annotation of 11 broadly defined populations of progenitor and differentiated cells from all germ layers that contribute to cardiac and pharyngeal arch development at multiple stages (Fig. 1a–1d and Extended Data Fig. 1b and 1c)^9^. Jaccard similarity analysis of cells between both modalities confirmed strong concordance of cell type annotation (Extended Data Fig. 1d). Peak calling in scATAC-seq identified 548,312 unique open chromatin regions across all cell types with 424,819 sites (77.5%) located in intronic and distal intergenic regions greater than 3 kB from known transcription start sites, indicating an enrichment for gene regulatory elements such as enhancers (Extended Data Fig. 1e and 1f). Differential accessibility analysis identified 205,037 cell type enriched chromatin regions, with each population containing open chromatin located distal to known cell type marker genes (Extended Data Fig. 1g). Motif analysis of cell type enriched chromatin regions revealed enrichment for lineage-specifying transcription factors critical for the development of each respective population (Extended Data Fig. 1h).

**Fig. 1.**
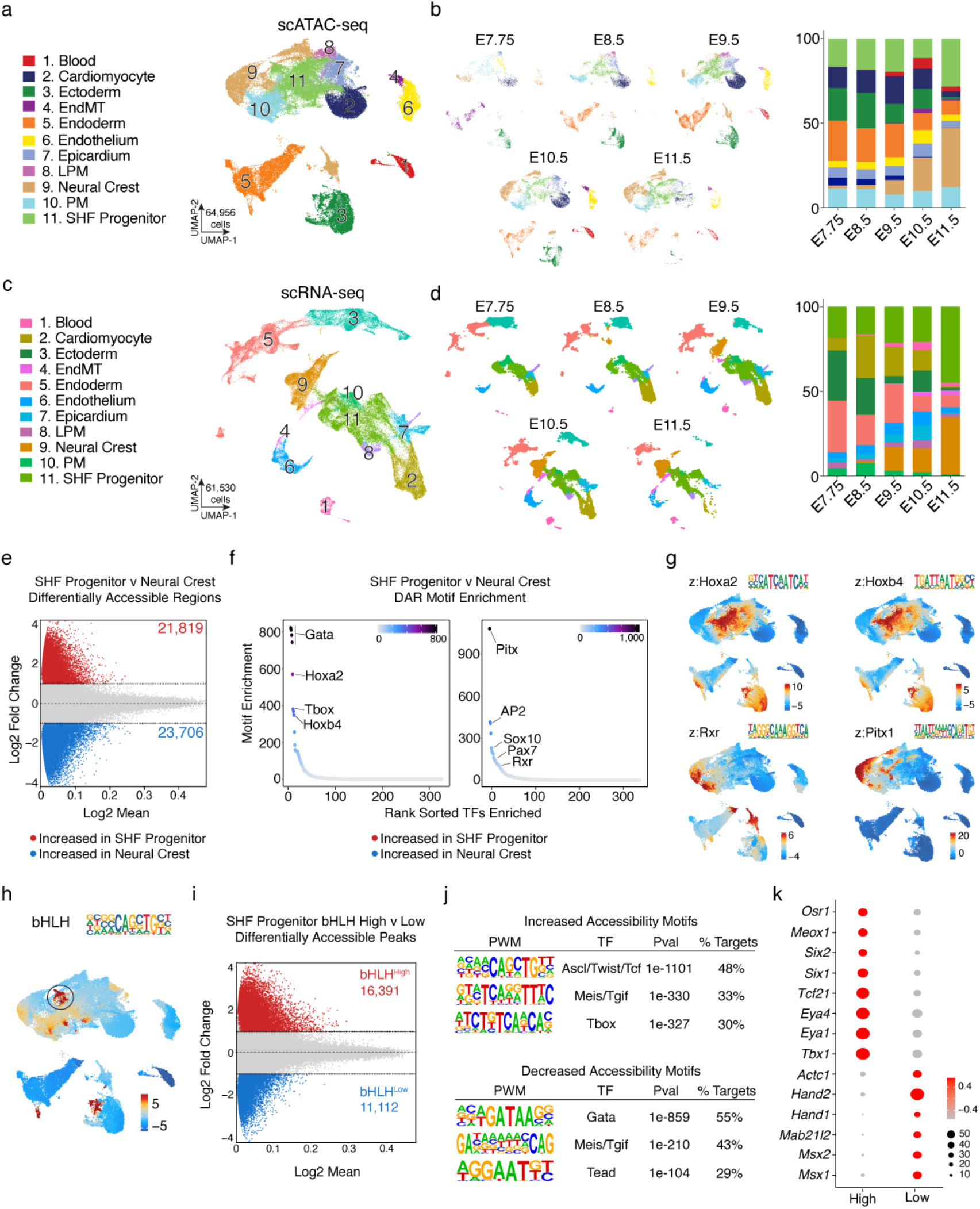
scATAC-seq Identifies cis-Regulatory Elements in Mouse Heart Development at Single Cell Resolution. **a,** Annotated UMAP of scATAC-seq aggregated data of 64,956 cells for cardiac crescent stage (E7.75, n=2), linear heart tube (E8.5, n=5), looped heart (E9.5, n=4), and chamber formation (E10.5, n=4) and early septation stages (E11.5, n =2). EndMT, Endothelial to Mesenchymal Transformation; PM, paraxial mesoderm; SHF Progenitor, second heart field progenitor; LPM, lateral plate mesoderm. Number of replicates represents individual embryos. **b,** UMAPs of scATAC-seq data separated by each time point (E7.75, n=4,002; E8.5, n=9,496; E9.5, n=20,035; E10.5, n=17,406; E11.5, n=14,017) and bar plots of cluster proportions per time point. Number of replicates represents cells. **c,** Annotated UMAP of scRNA-seq aggregated data of 61,530 cells for cardiac crescent stage (E7.75, n=5), linear heart tube (E8.5, n=2), looped heart (E9.5, n=4), and chamber formation (E10.5, n=4) and early septation stages (E11.5, n =2). Number of replicates represents individual embryos. **d,** UMAPs of scRNA-seq data separated by each time point (E7.75, n=10,483; E8.5, n=8,573; E9.5, n=18,261; E10.5, n=14,228; E11.5, n=10,021) and bar plots of cluster proportions per time point. Number of replicates represents cells. **e,** Differential accessibility testing (log FC >1 and FDR <0.05) of SHF Progenitor (n=12,291 cells) and Neural Crest (10,401 cells). Numbers in graph refer to number of differentially accessible sites that are increased (red) or decreased (blue) in accessibility in reference to SHF Progenitors. **f,** Plot of motifs enriched within sites increased or decreased in accessibility in SHF Progenitor cells. Rank sorted TF motifs (using HOMER) on x-axis and - log10 FDR of motif enrichment on y-axis. Scale bar represents mlog10 adjusted p-value range. **g,** UMAP of cells with motif enrichment for SHF-Progenitor and Neural Crest factors using ChromVAR and overlaid with motif logo from HOMER (scATAC). Scales for scATAC plot represent deviation scores, or z-scored values for each feature or motif, to vary from the average of all cells (colors are scaled where red represents the highest deviation for a given motif away from averages of all cells). **h,** UMAP of cells with motif enrichment for bHLH motif (matching Ascl1) using ChromVAR and overlaid with motif logo from HOMER (scATAC). Circled region depicts subcluster of interest within SHF-Progenitor population. **i,** Differential accessibility testing (log FC >1 and FDR <0.05) of bHLH-motif^High^ (n= 1568 cells) vs. bHLH-motif^Low^ (n=277 cells). Numbers in graph refer to number of differentially accessible sites that are increased (red) or decreased (blue) in accessibility in reference to SHF Progenitors. **j,** Top three enriched motifs from HOMER of differentially accessible regions increased in bHLH-motif^High^ and bHLH-motif^Low^ cells. **k,** Dot plot of differentially expressed genes in bHLH-motif^High^ vs. bHLH-motif^Low^ cells. Color intensity represents scaled expression levels and dot size is proportion of cells within cluster expressing the gene. Cells matched with Seurat’s canonical correlation analysis (CCA) were used for comparison.

Multipotent SHF-progenitor (#9) and neural crest cells (#11) signal reciprocally and differentiate to a wide range of derivative subpopulations, with cell fate decisions dictated by binding of lineage-specifying transcription factors to sequence-specific DNA motifs within *cis* regulatory elements^23^. Differential accessibility analysis comparing these two populations revealed thousands of unique regulatory elements (Fig. 1e). Gata and Pitx motifs displayed the highest enrichment in respective populations, in line with known roles for these factors in outflow tract formation^2,24^. Motifs for established differentiation factors such as Ap2, Sox10, and Pax7 were found in sites enriched in neural crest cells, whereas known transcription factors that regulate antero-posterior patterning, HoxA2 and HoxB4, were enriched in SHF-progenitors (Fig. 1f)^25,26^. To identify motifs enriched within subpopulations of SHF-progenitor and neural crest cells, we performed ChromVAR analysis on scATAC-seq peaks to calculate transcription factor motif deviation scores at single cell resolution (Fig 1g)^27^. Interestingly, ChromVAR revealed a distinct subpopulation of SHF-progenitor cells enriched for bHLH motifs and depleted for HoxA2 and HoxB4, marking cardiac progenitors (Fig. 1h)^28^. Subsetted analysis of this subpopulation (bHLH-motif^High^) compared to other SHF-progenitor cells (bHLH-motif^Low^) identified thousands of differentially accessible peaks with motifs enriched for pharyngeal muscle specification rather than cardiac determination (Fig. 1i and 1j). Integrated scRNA-seq analysis identified differential expression of numerous skeletal muscle-specifying transcription factors such as Meox1 and Six1/2 and decreased expression of cardiac lineage factors such as Hand1 and Hand2, highlighting the broad developmental potential of SHF progenitor cells^29^ (Fig. 1k). Overall, our scATAC-seq results define cis regulatory elements for all populations in early cardiogenesis and identify selective motif enrichment for developmentally essential factors within SHF and neural crest lineages.

### Computational Predictions of Heart and Neural Crest Enhancers

Although scATAC-seq can readily identify open chromatin at single cell resolution in heart development, accessibility alone is insufficient information to annotate the regulatory function of a genomic locus^30^. Amongst various types of *cis* regulatory elements, enhancers in particular play an essential role in controlling cell type restricted gene expression^10^. We therefore applied two methods to predict scATAC-seq peaks as likely enhancers, leveraging the integration of scRNA-seq and scATAC-seq data, as well as developing machine learning predictive models (Extended Data Fig. 2a). First, we used ArchR to identify linkages of scATAC-seq peaks within a 500kb boundary of a gene body locus and incorporated scRNA-seq values to determine which peaks may be linked to a transcribed gene^21^. This analysis not only predicts putative peaks as enhancers but also suggests the target gene, and led to 49,664 unique peak-to-gene linkages out of 548,312 total peaks (9%) (Table S2).

We also employed an orthogonal machine learning approach previously shown to have a high validation rate for predicting neurodevelopmental enhancers^31^. We first trained a supervised machine learning model to predict which scATAC-seq peaks are most similar to experimentally validated heart enhancers from the VISTA consortium (Fig. 2a)^32^. This model, agnostic to gene expression, predicts heart enhancer potential (positives) compared to regulatory potential in other tissues or a general lack of potential (negatives). The strategy enables high-confidence predictions using signatures of heart-specific enhancer activity, such as transcription factor binding, open chromatin or histone modifications in cardiac cells, rather than general signatures of enhancer activity common across all tissues. The trained model for prediction of heart enhancers uses 13 epigenetic features from mouse primary tissue and *in vitro* cardiac differentiations with varying weights (Extended Data Fig. 2b). Benchmarking analysis revealed the predictions from the machine learning model exceeded the performance of any individual feature or various combinations of ChIP-seq datasets in identifying validated heart enhancers (Extended Data Fig. 2c). The machine learning model was then used to score scATAC-seq peaks, with high-scoring peaks clustering near validated heart enhancers and low-scoring peaks clustering near non-heart enhancers (Extended Data Fig. 2d). Out of 548,312 total peaks, 25,114 peaks were confidently annotated as active heart enhancers (4.6% total, prediction scores >0.4). To identify putative lineage-restricted enhancers, we filtered the 237,674 cluster enriched peaks for sites with prediction scores > 0.4, leading to a considerably narrower set of 14,814 sites (2.7 % total peaks) (Fig. 2b). Peaks enriched within the cardiomyocyte cluster contained the largest number of predicted enhancers, whereas the ectoderm lineage contained the fewest, in line with model training that distinguishes heart from brain enhancers.

**Fig. 2.**
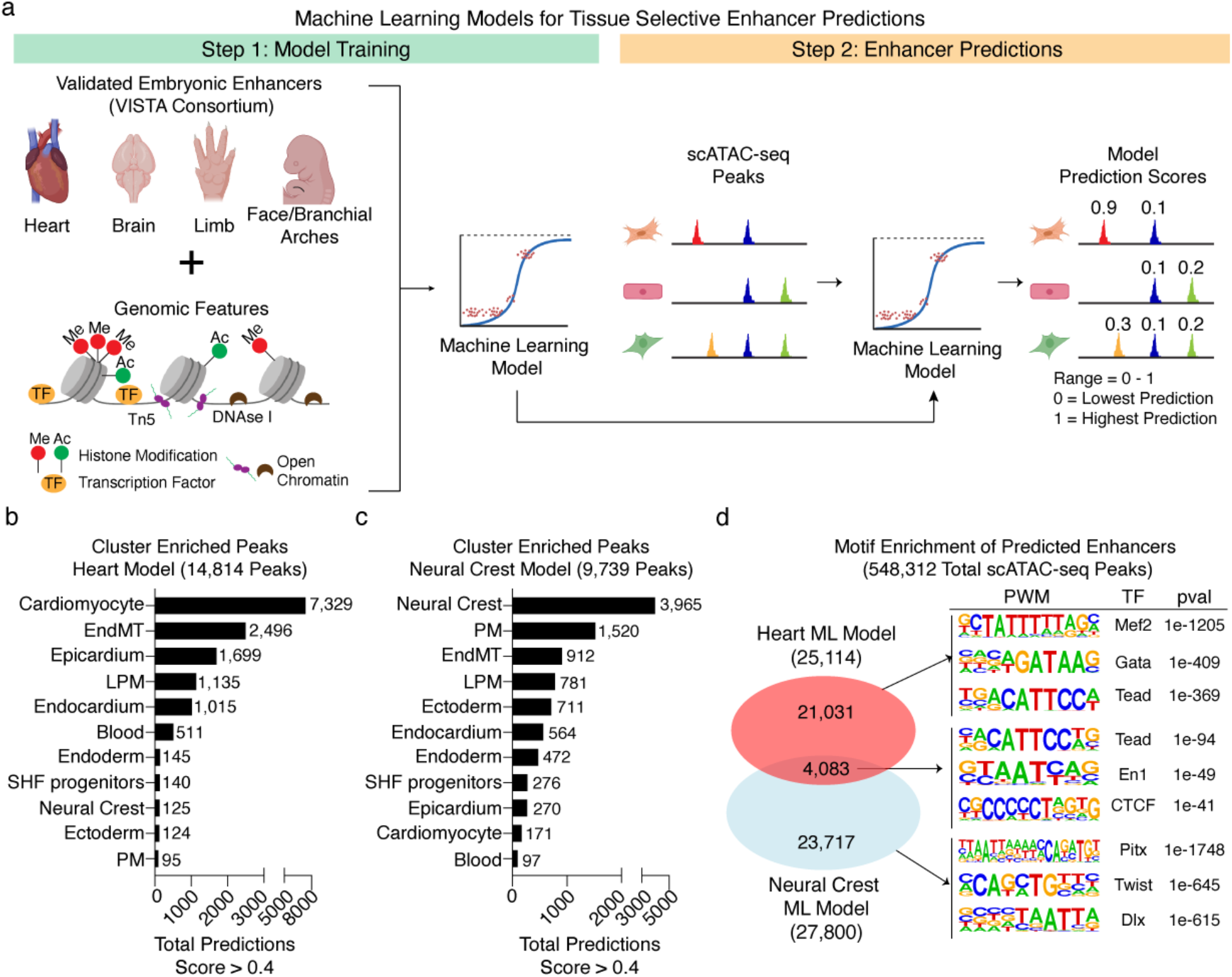
Machine Learning Models for Prediction of Heart and Neural Crest Enhancers. **a,** Cartoon illustration of machine learning model training and assignment of prediction scores to scATAC-seq peaks. In Step 1, a positive tissue type is selected using VISTA-validated embryonic enhancers annotated with expression within that population. Tissue types for the model to avoid (negative) are also chosen. Publicly available genomics datasets or features are then used for model training. In step 2, scATAC-seq peak coordinates are inputted into the trained model and each peak is assigned a prediction value score of 0 (unlikely to be a positive tissue type enhancer) to 1 (highly likely to be a positive tissue type enhancer). **b,** Bar plot of cluster-enriched peaks considered predicted heart enhancers (prediction score > 0.4) for each cluster using the heart enhancer trained model. **c,** Bar plot of cluster-enriched peaks considered predicted neural crest enhancers (prediction score > 0.4) for each cluster using the neural crest trained model. **d,** Venn diagram depicting overlaps between peaks considered enhancers within both models and motif enrichment of sites enriched within a respective model and overlapping regions using HOMER.

Notably, of the 16,361 neural crest cell-enriched peaks, the heart enhancer model identified very few as putative enhancers (125/16,361 or 0.76%). To improve neural crest enhancer predictions, we generated another machine learning model following a similar approach as described above but using VISTA enhancers assayed in tissues relevant to neurocristopathies, such as facial mesenchyme, branchial arch and mesenchyme derived from neural crest^32^. This new model uses 37 epigenetic features from mouse primary tissue and dramatically increased our ability to identify neural crest enhancers (3,965/16,361 or 24%), highlighting the flexible nature of model training to define cell type enriched regulatory elements (Fig. 2c and Extended Data Fig. 2e). Interestingly, the neural crest model also identified more enhancers in paraxial mesoderm cells (1,520/13,403 or ~0.5%), which give rise to mesenchymal and skeletal derivatives. Intersection of predicted enhancers from both models revealed 2,628 sites that contained not only known motifs for cardiac (Tead) and neural crest progenitor cells (En1), but also CTCF, an essential DNA-binding factor with roles in structural genome organization and general transcriptional regulation (Fig. 2d)^33^.

We then experimentally tested predicted enhancers from both machine learning models *in vivo* for regulatory activity (Table S3). For the heart enhancer model, we cloned 5 novel peaks with high cardiomyocyte enhancer prediction and high conservation into a Tol2-GFP reporter vector and injected the reporters into F_0_ cmlc2:mCherry Zebrafish (D. *rerio*) embryos, which marks cardiomyocytes. Examination of GFP expression at 72 hours post fertilization (hpf) revealed enhancer activity in the expected cell type for 4 out of 5 sites despite the cross-species distance, and we present examples for 3 representative enhancers (Extended Data Fig. 3a-f). Notably, only one of the three predicted enhancers was situated within an intron of a gene expressed in cardiomyocytes (*Sh3bgr*), whereas the other two peaks were either within an intron of a gene ubiquitously expressed or distal to a gene not expressed at all (Extended Data Fig. 3g). For the neural crest enhancer model, we cloned 8 sites with high prediction scores into a Tol2-GFP reporter vector and injected them into F0 Tg(sox10-mRFP) Zebrafish embryos, in which neural crest cells are marked. We identified overlapping GFP expression with mRFP from all 8 regions tested (8/8, 100%) and present examples for 3 representative enhancers (Extended Data Fig. 4a-f). Expression analysis of nearest genes again failed to identify neural crest-specific expression, suggesting the machine learning approach is able to identify potentially long-rage enhancers regulating nonneighboring genes (Extended Data Fig. 4g). These results highlight the unique capability of machine learning approaches to uncover novel cell type specific regulatory elements without *a priori* knowledge of proximal gene expression.

Overall, our integrated scATAC-seq dataset and computational cell type enhancer prediction approaches provide an unprecedented resource of the spatiotemporal *cis* regulatory landscape during mouse heart development at single cell resolution. The knowledge of cellular expression of secreted proteins and their cognate receptor expression in neighboring cells, along with downstream chromatin consequences in signal receiving cells, forms a foundation for mechanistic investigation of non-cell autonomous consequences of cellular communication during development and disease.

### Tbx1 Deficiency Broadly Disrupts Cell Signaling with Posteriorization of Anterior Second Heart Field Progenitors

Having established a framework to interrogate epigenetic and transcriptomic events during heart development at single cell resolution,–we leveraged this knowledge to address a long-standing question relevant to human disease. Since TBX1 functions non-cell autonomously in SHF cells to affect neural crest cells leading to cardiac and craniofacial defects in 22q11 deletion syndrome^20^, we performed integrated scATAC-seq and scRNA-seq analysis of WT and Tbx1-null embryos between E9.5 - E11.5, micro-dissecting regions enriched for SHF-progenitors and neural crest cells (Extended Data Fig. 5a). Because deletion of Tbx1 in mesoderm-derived lineages is necessary and sufficient for cardiac defects, we focused our analysis on SHF-derived mesodermal progenitors and neural crest populations^34^. We applied similar analyses for scRNA-seq and scATAC-seq as previously described and annotated 10 cell types based on expression and gene score plots of known marker genes, respectively (Fig. 3a-3d and Extended Data Fig. 5b and 5c). Integration of data from both modalities led to high concordance of cell type annotation (Extended Data Fig. 5d). Notably, the proportions of cell types were similar between genotypes at all time points except neural crest cells at E11.5, where we noticed a pronounced reduction in Tbx1-null embryos in both modalities (FDR <0.05, Fig. 3b and 3d).

**Fig. 3.**
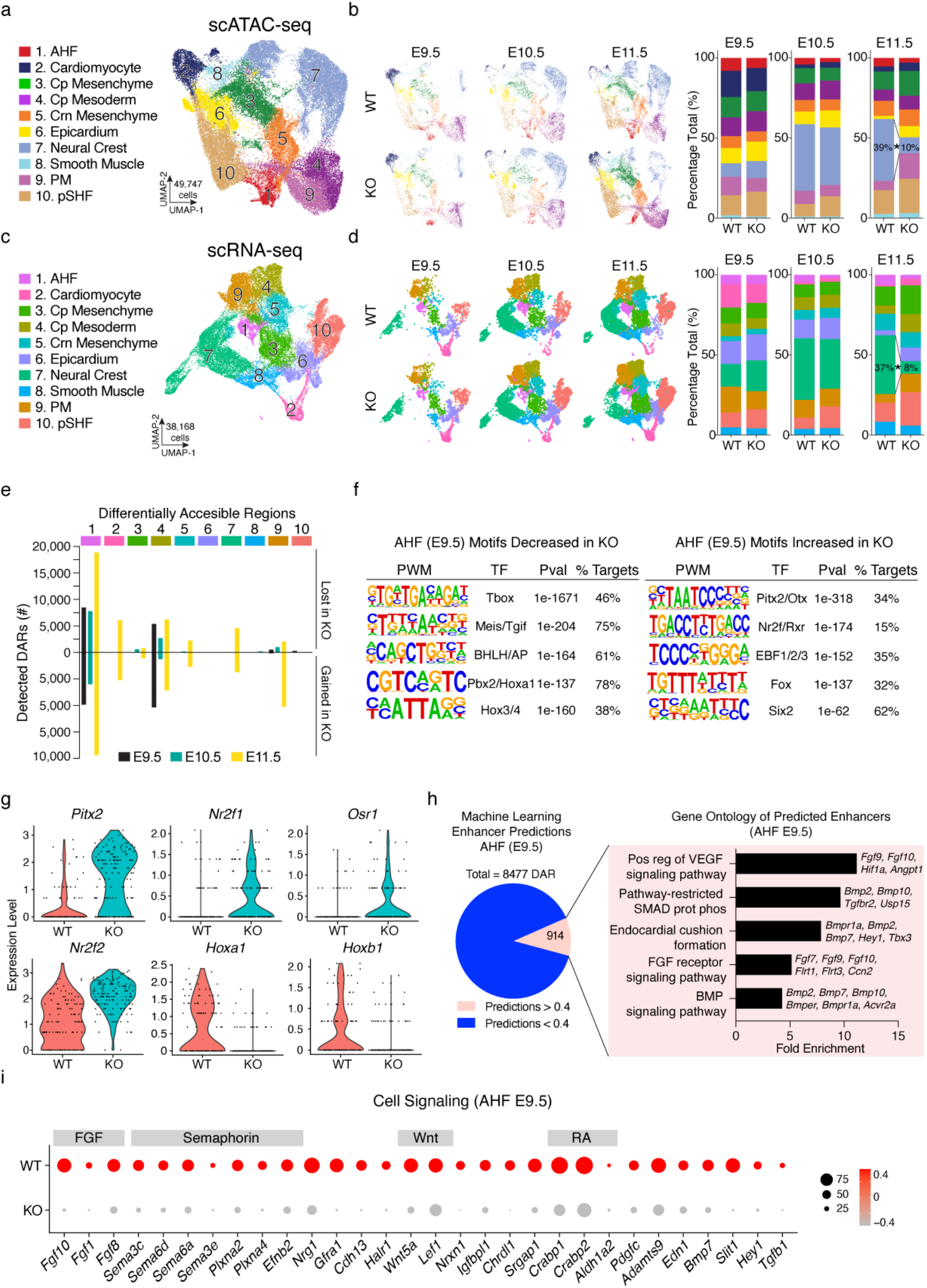
Loss of Tbx1 Disrupts Expression of Cell Signaling Genes and Anterior-Posterior Patterning in SHF. **a,** Annotated UMAP of scATAC-seq cells within mesoderm lineage and neural crest cells (n = 49,747 cells) including both WT and KO cells at E9.5 (n= 2 WT and 2 KO reps), E10.5 (n=2 WT and 2 KO reps) and E11.5 (n=2 WT and 3 KO reps). AHF, anterior heart field; pSHF, posterior second heart field; CPM, cardiopharyngeal mesoderm; PM, paraxial mesoderm; Cr Mes, cranial mesenchyme; Cp Mes, cardiopharyngeal mesenchyme. **b,** UMAP images and bar plot of samples for WT (E9.5, n=4,806; E10.5, n=7,120; E11.5, n=11,150 cells) and KO (E9.5, n=4,519; E10.5, n=7,339; E11.5, n=14,813 cells). Percentages for neural crest are shown at E11.5 to highlight differences between WT and KO. Statistical test conducted with permutation test, FDR < 0.05 and abs(log2FD) > 0.58. **c,** Annotated UMAP of scRNA-seq cells within mesoderm lineage and neural crest cells (n = 38,168 cells) including both WT and KO cells at E9.5 (n= 2 WT and 2 KO reps), E10.5 (n=2 WT and 2 KO reps) and E11.5 (n=2 WT and 3 KO reps). **d,** UMAP images and bar plot of samples for WT (E9.5, n=1,858; E10.5, n=5,048; E11.5, n=8,592 cells) and KO (E9.5, n=2,500; E10.5, n=6,531; E11.5, n=13,639 cells). Percentages for neural crest are shown at E11.5 to highlight differences between WT and KO. Statistical test conducted with permutation test, FDR < 0.05 and abs(log2FD) > 0.58. **e,** Differential accessibility testing (log FC >1 and FDR <0.05) of Tbx1^+/+^ and Tbx1^-/-^ cells in mesoderm derived clusters at three time points. Color code and numbers refer to scATAC-seq cell type identities from panel a and legend of colors refer to time points. **f,** Top five motifs enriched in sites lost and gained within AHF cells at E9.5 using HOMER. **g,** Violin plot of retinoic acid signaling dependent factors normally expressed in posterior second heart field within AHF cells at E9.5 using Wilcoxon test (logfc.threshold = 0.25 and min.pct = 0.1). **h,** Venn diagram of sites lost in AHF cells at E9.5 filtered by sites considered predicted enhancers using the cardiac linear model (prediction scores > 0.4) and bar plot of fold enrichment for signaling pathways affected in AHF cells at E9.5 using Gene Ontology analysis (Panther 17.0 Overrepresentation Test) of 914 sites considered putative enhancers. **i,** Dot plot of differentially expressed cell signaling genes in AHF cells at E9.5. Color intensity represents scaled expression levels and dot size is proportion of cells within cluster expressing the gene.

To determine which populations were affected by loss of Tbx1, we first performed differential accessibility testing comparing genotypes within each cell type at all three time points and identified aberrant chromatin accessibility in discrete subpopulations (Fig. 3e). The SHF is subdivided into a Tbx1-high anterior compartment that contributes to the anterior heart field (AHF) (cluster 1) and cardiopharyngeal mesoderm progenitors (cluster 4), and a Tbx1-low posterior SHF (pSHF) (Extended Data Fig. 5e). Chromatin of the AHF and cardiopharyngeal mesoderm were the most affected at all time points upon loss of Tbx1 (Fig. 3e). Other populations such as the paraxial mesoderm (cluster 9) express Tbx1 at all time points tested but displayed noticeable epigenomic alterations only at E11.5 (Fig. 3e and Extended Data Fig. 5e). Given that Tbx1 deficiency leads to outflow tract anomalies, and because AHF progenitors give rise to outflow tract cardiomyocytes and communicate with migrating neural crest cells, we focused our analysis on this population at the earliest time point, E9.5. Motif analysis among regions with decreased accessibility within AHF cells at E9.5 identified a Tbox motif as the most enriched (~50% of peaks), suggesting regions directly affected by loss of Tbx1 (Fig. 3f). Motifs for AHF transcription factors required for antero-posterior patterning including Isl1, Hoxa2 and Meis, previously implicated in SHF and neural crest patterning, were also present in closed chromatin regions^28^. Amongst sites with increased accessibility, loss of Tbx1 led to enrichment of motifs for the segmentation and symmetry patterning transcription factor Pitx2^35^. Dysregulated regions also contained motifs for neural transcription factors Otx2 and Ebf2, in line with a previously suggested role for Tbx1 to suppress neural fate (Fig. 3f)^36^. Notably, ectopically accessible regions contained motifs for Nr2f, Foxf and Rxr transcription factors that typically drive the pSHF fate, giving rise to atrial myocardial cells (Fig. 3f)^37^. Concordantly, scRNA-seq analysis revealed decreased expression of Hoxa1 and Hoxb1 along with an increase in expression of atrial factors Nr2f1, Nr2f2 and Osr1 as well as Pitx2, essential for left/right patterning and regulated by retinoic acid signaling (Fig. 3g)^38^. Although Tbx1 had previously been shown to coordinate a boundary of anterior and posterior heart fields, our data suggest a novel function of aberrant reprogramming of AHF cells, failing to suppress non-mesoderm lineages, and adopting more pSHF characteristics upon loss of Tbx1^39,40^.

To identify putative enhancers amongst the dysregulated sites within E9.5 AHF progenitors, we applied our heart enhancer machine learning model to sites lost in Tbx1 KO cells and identified 914/8477 (~11%) predicted enhancers (Fig. 3h). Gene ontology results of genes distal to the 914 predicted enhancers revealed an enrichment for cell signaling terms including FGF, VEGF and BMP pathways (Fig. 3h). Differential gene expression results from scRNA-seq analysis revealed a dramatic decrease in numerous signaling pathways required for outflow tract development including FGF, Semaphorin, Wnt, BMP and Notch genes, suggesting that Tbx1 is a master regulator of gene programs for cell signaling in AHF cells (Fig. 3i)^41^.

To correlate the changes in gene expression to Tbx1-dependent chromatin accessibility changes, we intersected differentially accessible peaks and expressed genes and found 257 downregulated genes distal to decreased scATAC-seq peaks containing a Tbox binding site, suggesting sites directly regulated by Tbx1 (Extended Data Fig. 6a). Amongst these dysregulated genes, Sema3c disruption, similar to loss of Tbx1, leads to outflow tract anomalies such as common arterial trunk^42^. We identified two scATAC-seq peaks in AHF cells distal to Sema3c that were lost in Tbx1 KO cells and predicted to contain a Tbox motif (Extended Data Fig. 6b). We cloned both sequences into a luciferase reporter assay vector and found that both sites displayed robust activity when cotransfected with a mouse *Tbx1* cDNA vector. Deletion of the 10bp putative Tbox site led to a significant decrease in activity, suggesting that both sites are likely Tbx1-dependent enhancers for Sema3c in AHF cells (Extended Data Fig. 6c).

Although AHF progenitor cells differentiate to outflow tract and right ventricular myocardium, scATAC-seq revealed differential chromatin accessibility only in later stage cardiomyocytes (E11.5) but not earlier stages (E9.5 and E10.5) (Extended Data Fig. 7a). In line with differential motifs indicating alterations to antero-posterior patterning in early-stage AHF cells, a subsetted analysis of outflow tract cardiomyocytes at E11.5 revealed a marked increase in expression of atrial myocardial genes such as Tbx5, Nr2f1, Nr2f2 and Wnt2 and downregulation of an outflow tract myocardial signature of Rspo3, Bmp4 and Sema3c and Isl1 (Extended Data Fig. 7b)^9^. Importantly, to rule out the possibility that outflow tract and ventricular myocardium displayed specification defects in stages prior to our analysis, we performed scATAC-seq at E8.25 (linear heart tube stage) in WT and Tbx1-deficient embryos (Extended Data Fig. 7c and 7d). After performing quality control, clustering and subsetting for mesoderm lineage populations as previously described, we annotated 10 cell types, based on gene score plots of known marker genes, that included SHF progenitors and differentiated myocardial subtypes (Extended Data Fig. 7e). Equivalent numbers of Tbx1 WT and KO cells were again found in each population, indicating that loss of Tbx1 did not affect the initial differentiation of these cell types (Extended Data Fig. 7d). Similar to E9.5, the chromatin landscape of AHF cells was dramatically altered, with closed regions enriched in cardiac factors Tbox, Isl1 and Pbx motifs and ectopically gained sites containing motifs for pSHF specification and retinoic acid metabolism such as Nr2f, Pitx2, Foxf and Rxr factors (Extended Data Figs. 7f and 7g). No appreciable defects in chromatin accessibility were identified in outflow tract and ventricular myocardium, suggesting Tbx1 is dispensable for the initial specification of these differentiated populations, but rather functions primarily in cell signaling and in the anterior-posterior patterning of the AHF (Extended Data Fig. 7f).

### Axial Patterning Defects in Cardiopharyngeal Neural Crest Cells in Response to Tbx1-Dependent Signaling

We next investigated the non-cell autonomous effects due to loss of Tbx1 on the epigenetic and transcriptomic features of neighboring neural crest cells, which have negligible Tbx1 expression (Extended Data Fig. 5e). To resolve the heterogenous populations of neural crest cells captured in our analysis, we selected and re-clustered cells based on expression of well-established neural crest transcription factors Twist1 and Dlx2/5, leading to four broad populations of cells in both modalities (Figs. 4a-4d, Extended Data Figs. 8a and 8b)^3^. scRNA-seq clearly identified restricted gene expression profiles for each population, and these marker genes were also segregated at the level of chromatin accessibility in gene score plots (Extended Data Figs. 8a and 8b). The transcription factors Sox10 and Foxd3 were expressed uniquely within the migratory multipotent progenitor cells (Progenitor NC) whereas craniofacial NC cells expressed Ebf1 and Smoc1, transcription factors essential for cranial muscle and mesenchyme derivatives^43,44^. Interestingly, we identified two distinct populations of cardiac neural crest derivatives: a more differentiated cardiac outflow valve cushion population marked by transcription factors Isl1, Hand1 and Hand2 (Cardiac NC), as well as an independent cluster expressing high levels of progenitor-enriched factors Hox3 and Hox4, known to be expressed in the caudal pharyngeal arches (PA3-6) where cardiac neural crest cells migrate^13^. RNAscope-based in situ hybridization confirmed the Hoxb3-expressing neural crest derivatives were localized mainly to the region dorsal to the outflow tract around PA4 and likely represent a progenitor cardiac neural crest population distinct from the craniofacial neural crest cells, in line with clustering analysis (Extended Data Figs. 8c and 8d). Each of the four neural crest subsets contained thousands of unique accessibility sites distal to population marker genes and were enriched for motifs of distinct lineage-specifying factors for cranial (e.g. Pitx, Brn1 and Olig2) or cardiac (e.g. Fox and Gata6) neural crest cells (Extended Data Figs. 8e and 8f). Analysis of cell proportions between WT and KO revealed no differences in early stages of pharyngeal neural crest populations (E9.5 and E10.5), but did identify a marked decrease of craniofacial NCs at E11.5, along with a concordant expansion of progenitor cells, suggesting differentiation defects (Figs. 4b and 4d).

**Figure 4.**
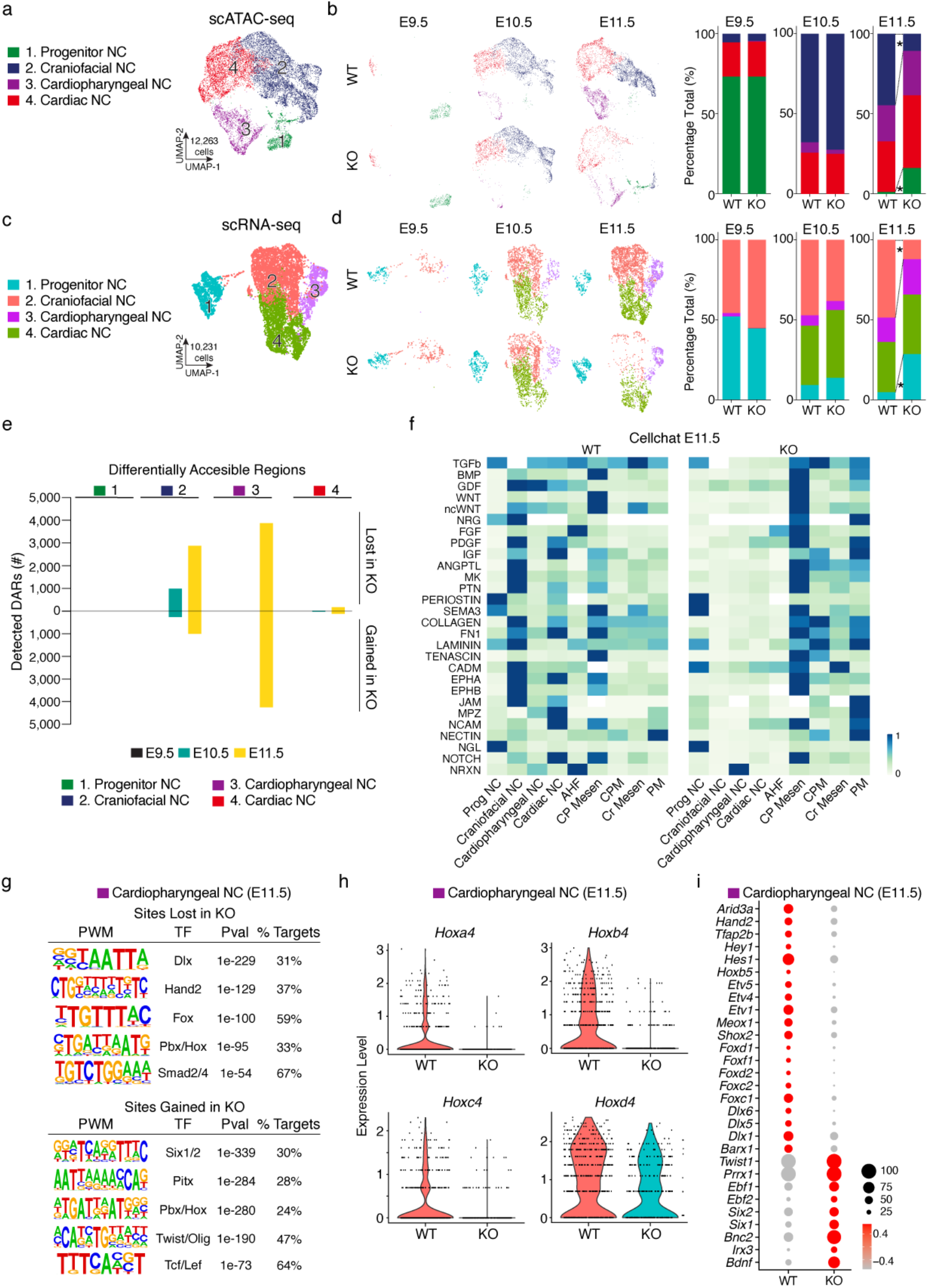
Axial Patterning and Differentiation Defects in Cardiac Neural Crest Population. **a,** Annotated UMAP of scATAC-seq cells within neural crest subset (n = 12,263 cells) including both WT and KO cells at E9.5 (n= 2 WT and 2 KO reps), E10.5 (n=2 WT and 2 KO reps) and E11.5 (n=2 WT and 3 KO reps). NC, neural crest. **b**, UMAP images and bar plot of samples for WT (E9.5, n=421; E10.5, n=2,925; E11.5, n=4,305 cells) and KO (E9.5, n=503; E10.5, n=2,616; E11.5, n=1,493 cells). Stars indicate significant differences between E11.5 WT and KO in Progenitor NC (1% WT, 16% KO) and Craniofacial NC (45% WT, 11% KO). Statistical test conducted with permutation test, FDR < 0.05 and abs(log2FD) > 0.58. **c,** Annotated UMAP of scRNA-seq cells within neural crest subset (n = 10,231 cells) including both WT and KO cells at E9.5 (n= 2 WT and 2 KO reps), E10.5 (n=2 WT and 2 KO reps) and E11.5 (n=2 WT and 3 KO reps). **d,** UMAP images and bar plot of samples for WT (E9.5, n=286; E10.5, n=2,109; E11.5, n=3,703 cells) and KO (E9.5, n=507; E10.5, n=2,208; E11.5, n=1,418 cells). Stars indicate significant differences between E11.5 WT and KO in Progenitor NC (5% WT, 29% KO) and Craniofacial NC (49% WT, 12% KO). Statistical test conducted with permutation test, FDR < 0.05 and abs(log2FD) > 0.58. **e,** Differential accessibility testing (log FC >1 and FDR <0.05) of Tbx1^+/+^ and Tbx1^-/-^ cells in neural crest subpopulations at E9.5, E10.5 and E11.5. Color code and numbers refer to scATAC-seq cell type identities from panel a and legend of colors refer to time points. **f,** Heatmap of overall signaling patterns (both incoming and outgoing) for four neural crest cell types and mesoderm lineage populations for both WT and KO cells at E11.5. Relative strength values are z-scores values of overall signaling patterns for curated set of statistically significant pathways, as assessed by Cellchat. **g,** Motif enrichment analysis of sites lost and gained in KO within cardiopharyngeal neural crest cells at E11.5 using HOMER. **h,** Violin plot of log normalized values for Hox4 cluster genes within the cardiopharyngeal neural crest population at E11.5. **i,** Dot plot of differentially expressed genes in cardiopharyngeal neural crest cells at E11.5. Color intensity represents scaled expression levels and dot size is proportion of cells within cluster expressing the gene.

Having refined the neural crest subpopulations, we first asked which incoming signaling pathways were important for each of the four subpopulations. Cellchat network signaling analysis revealed distinct pathways enriched in each population (Extended Data Fig. 9a). Cardiac and craniofacial derivates shared overlapping connections (pattern 1) with BMP, Wnt, non-canonical Wnt and FGF signaling, whereas the cardiopharyngeal population (pattern 3) included members of the VCAM, Tgfb and Endothelin (Edn) pathways^45^. The migrating progenitor cells were linked to a unique subset (pattern 4) that included EGF and NRG signaling, known to be critical for trunk neural crest migration^46^.

We then performed differential accessibility testing for each subpopulation, comparing WT and KO at all time points, and identified aberrant chromatin accessibility predominantly within the cardiopharyngeal and craniofacial cell types at E11.5 (Fig. 4e). Although previous studies had implicated impaired migration of neural crest cells due to loss of Tbx1, our scATAC-seq results identified no epigenomic changes in the progenitor neural crest population at all three time points, indicating that the initial migratory multipotent cells were not affected due to loss of Tbx1-dependent signaling^16^. We next sought to define the disrupted cell signaling pathways between these cells and specific SHF progenitors affected by loss of Tbx1. We noted pronounced differences in signaling between neural crest and mesoderm progenitors at E11.5 (Fig. 4f). Numerous pathways known to affect neural crest lineages in the pharyngeal arches such as Tgfb, BMP, FGF and ncWNT displayed decreased overall signaling strength. Semaphorin, Notch and Ephrin pathways, which coordinate the direction and migration of neural crest cells, were also notably decreased. Focusing on the cardiopharyngeal neural crest DARs, we filtered dysregulated sites using both the cardiac and neural crest machine learning models to identify putative enhancers (Extended Data Fig. 9b). Amongst the regions ectopically activated, we noticed an increase in the proportion of sites considered neural crest and not heart enhancers, suggesting that loss of Tbx1-dependent signaling may be causing failed repression of neural crest progenitor identity in this cardiac fated population (Extended Data Fig. 9b).

Differential accessibility of the cardiopharyngeal neural crest cells at E11.5 revealed thousands of dysregulated sites (Fig. 4e). Among aberrantly closed regions, there was motif enrichment for transcription factors involved in neural crest differentiation such as Dlx members, as well as cardiac neural crest differentiation factors such as Hand2, Fox and Hox (Fig. 4g). Consistent with this, sites that were ectopically accessible in the Tbx1 mutants were enriched for motifs that bind transcription factors involved in craniofacial and pharyngeal mesenchyme development such as Six1/2, Pitx and Twist. This suggests that loss of Tbx1 signaling may impair the repression of a network of genes typical of more rostral pharyngeal arches, involved in craniofacial development, and result in failure of the more caudal arches involved in cardiac development to activate their normal gene program.

In line with axial patterning defects found in AHF progenitors, scRNA-seq of cardiopharyngeal neural crest cells revealed decreased expression of numerous Hox4 factors, providing a mechanistic link underlying previously described hypoplasia of the fourth pharyngeal arch in Tbx1 null embryos (Fig. 4h)^34^. Differential gene expression revealed decreased levels of numerous transcription factors critical for cardiac neural crest lineages including Hand2, Tfap2b, Etv and Fox factors (Fig. 4i). Amongst genes with increased expression levels, multiple factors normally restricted to craniofacial NCs such as Ebf1/2, Prrx1 and Twist1 were ectopically high, indicating a failed repression of cranial identity. Finally, we intersected the machine learning predicted enhancers with decreased accessibility with differentially expressed genes and identified two putative enhancers distal to the FGF responsive genes Spry1 and Spry2, which were both decreased in expression in Cardiopharyngeal NCs at E11.5 and previously shown to genetically interact with Tbx1 (Extended Data Figs. 9c-f)^47^. Thus, the neural crest defects in Tbx1 mutants appear to reflect a failure of proper anterior-posterior patterning of discrete neural crest-derived cells in response to dysregulated anterior-posterior organization of the SHF.

## Discussion

Communication between diverse cell types within the cardiac outflow tract is essential for proper morphogenesis, and alterations in this process are a main cause of congenital heart defects^48^. Here, we employed single cell multimodal analyses to comprehensively define the transcriptional states and regulatory networks of the diverse cells involved in morphogenetic steps during cardiogenesis and reveal discrete defects within specific subpopulations of cells affected by loss of Tbx1. Previous studies have shown the requirement of Tbx1 in regulating FGF8 and other signaling factors in cardiac development, however the cell types involved and mechanisms underlying downstream neural crest differentiation defects within the pharyngeal arches were poorly understood^49,50^. Our results suggest that loss of Tbx1-dependent signals in AHF cells coincides with impaired retinoic acid signaling, aberrant axial patterning and an increase in pSHF signatures. Increased expression of NR2F and FOXF factors and decreased expression of Hox genes correlated with increased expression of Pitx2, a key driver of left/right asymmetry in development^38^. Importantly, our single cell analyses identified two discrete populations of cardiac neural crest cells affected non-cell autonomously by loss of Tbx1 and defined differentiation defects within a subset of cells enriched for Hoxb3, a population previously shown to be regulated by retinoic acid signaling^51^. Within the affected cardiac neural crest population, decreased expression of Hox4 cluster genes, typical of PA3-6, correlated with increased expression of craniofacial factors typical of PA1 and PA2, such as Six1/2, Ebf1/2, Twist1 and Prrx1, along with the presence of motifs for these factors within ectopically accessible sites. Thus, our results suggest a model where the “anteriorization” of the cardiopharyngeal arches secondary to the “posteriorization” of the AHF underlies the morphogenetic defects associated with loss of TBX1 in 22q11 deletion syndrome. Future studies should uncover mechanistic links between changes to Pitx signaling and left/right symmetry defects in outflow tract and aortic arch remodeling, often seen in 22q11 deletion syndrome. Our scATAC-seq results and enhancer predictions offer novel insights into the regulatory elements driving these specific neural crest derivatives and allow for the generation of new reporter tools to perturb these cells. We hypothesize that aberrant differentiation and migration of this cardiopharyngeal population may underlie the varying penetrance and expressivity of outflow tract defects in humans who are haploinsufficient for Tbx1 in 22q11 deletion syndrome.

Our work demonstrates that in order to understand how cells cooperate to shape organs it is essential to have a temporal single cell map of the concurrent epigenetic and transcriptomic state of each cell. Here, we provide such a map over a critical time frame in cardiac morphogenesis when progenitor cells migrate into the cardiac region. We employ a machine learning approach to predict enhancers for both cardiac and neural crest lineages and use these predictions to filter differentially accessible chromatin peaks for functionality. We anticipate that future refinement of enhancer prediction models using CRISPR perturbations with single cell readouts will greatly improve the accuracy of model predictions and could ultimately identify novel regulatory elements and genes in cardiac and craniofacial development. A similar approach for each organ will ultimately reveal mechanisms underlying myriad birth defects at a depth previously unachievable. Overall, our work highlights the power of integrated single cell genomics to decipher the complex inter- and intra-cellular regulatory networks during organ formation in development and in disease.

## Methods

### Animal Models

All animals used in this study were handled in accordance to protocols approved by the Institutional Animal Care and Use Committees (IACUC) at the University of California San Francisco. All protocols were conducted in accordance to guidelines from the National Institutes of Health Guide for the Care and Use of Laboratory Animals. Wildtype C57Bl6/J mice (Stock:000664) were obtained from the Jackson Laboratory (Bar Harbor, ME). Tbx1 null mice were generated by breeding Tbx1 flx mice to E2A-Cre (Jackson Laboratory, Stock:003724)^1^. The Cre allele was outcrossed and Tbx1 heterozygous mice were maintained on C57Bl6/J background for > 10 generations.

All zebrafish (*Danio rerio*) husbandry was performed following UCSF guidelines under standard conditions. All experiments were conducted with larvae between 48 and 72hpf. The following transgenic lines were used in this study: outbred Ekkwill (EKW), Tg(cmlc2: mCherry)^2^ and Tg(*sox10:mRFP*)^3^.

### Embryo Harvest and Genotyping

For all timed mating experiments, male and female mice were housed overnight and female mice were visually inspected for a vaginal plug in the morning. Embryos at date of plug were considered E0.5. Females were checked for pregnancy at E6.5 by echocardiography (Vevo 3100, Visual Sonics). Embryos were harvested at indicated time points by euthanizing pregnant females according to approved protocols and collecting embryos in ice cold PBS-F, PBS (Life Technologies, 14190250) and 1% Fetal Bovine Serum on ice (Thermo Fisher Scientific, 10439016). Experiments for C57Bl6/J embryos did not require genotyping whereas, for Tbx1 embryos, yolk sacs were dissected and a small fraction was placed in QuickExtract DNA Extraction Solution (Lucigen, QE09050) and processed according to manufacturer’s protocols.

Tbx1 genotyping was performed by PCR using Phire Green Hot Start II DNA Polymerase (Thermo Fisher Scientific, F124L), according to manufacturer’s protocols, using primers to detect WT (P1+P2) and KO alleles (P1+P3):

P1: CGTTCTGTGCCTCTGACTGCGTGTC
P2: TCTACTGGTCAAATCGGGAGCATCTG
P3: GGTACTTGTAGGGATGGTGGTGCAGC

### Embryo Microdissection for scRNA-seq and scATAC-seq

WT C57Bl6/J embryos were selected by somite count stages: E7.75 (~4s), E8.5 (6-8s), E9.5 (~20-24s) and E10.5 (~32-35s). Stage-matched Tbx1 WT and KO embryos were used at E9.5, E10.5 and E11.5 (~42-45s). For embryos < E9.5, the entire cardiogenic region including the regions dorsal to the heart proper and including the pharyngeal arches were included. For embryos at E10.5 and E11.5, the regions dorsal to the heart were emphasized and minimal regions of the heart, specifically the outflow tract, was taken. At E11.5, the first pharyngeal arch was excluded to maximize the contribution of pharyngeal arches 3-6. Micro-dissected tissue was dissociated using either TrypLE Select (Thermo Fischer Scientific, 12563011) or Liberase TL (0.25 mg/ml in PBS, Millipore Sigma, 5401020001) for 10 minutes at 37C. Dissociated cells were quenched with PBS-F, strained through a 35-μm cell strainer (BD Falcon, 08-771-23) and centrifuged at 150 x g for 3 minutes. The cell pellet was resuspended in PBSF and a small aliquot was used for manual cell counting (SKC hemocytometer, 22-600-101, Fisher Scientific). For scRNA-seq experiments, a maximum of 10,000 cells per sample were used for scRNA-seq library preparation. For scATAC-seq experiments, the remaining volume of cells was again centrifuged at 150 x g for 3 minutes and resuspended in 100 ul ice cold nuclei lysis buffer for 5 minutes at 4 C (10 mM Tris-HCl, pH 7.4, 10 mM NaCl, 3 mM MgCl_2_, 0.1% Tween-20, 0.1% Nonidet P40 Substitute, 0.01% Digitonin and 1% Bovine Serum Albumin). After lysis, 500 ul wash buffer (10 mM Tris-HCl, pH7.4, 10 mM NaCl, 3 mM MgCl_2_, 0.1% Tween-20 and 1% BSA) was added and centrifuged at 500 x g for 5 minutes at 4 C. Nuclei were resuspended using 1X Nuclei Buffer (10x Genomics, PN-2000153/2000207). A maximum of 10,000 nuclei per sample in 5 ul volume were used for scATAC-seq preparation.

### scRNA-seq and scATAC-seq Sample Library Preparation

For scRNA-seq library preparation, cells were loaded onto the 10X Genomics Chromium instrument according to manufacturer’s protocols (Chip G, PN-1000120). All experiments were conducted with version 3.1 NEXT GEM reagents (PN-1000121), using 9 cycles of cDNA amplification for the GEM kit (PN-1000123) and 9 cycles of library amplification for library kit (PN-1000157). Each sample was indexed with a unique sample primer (Single Index Kit T Set A, PN-000213). For scATAC-seq library preparation, nuclei were first transposed with Tn5 enzyme (PN-2000138) and then loaded onto the Chromium instrument (Chip H, PN-1000161) using the version 1.1 NEXT GEM reagents (PN-1000175). Library preparation proceeded with 12 cycles each of GEM barcoding and library amplification. Each sample was indexed with a unique sample primer (Single Index Plate N Set A, PN-3000427). For both scRNA-seq and scATAC-seq, samples were pooled separately into respective libraries and sequenced on an Illumina NextSeq 500 (Illumina, software 4.0.2) and/or a lane of a Novaseq S4 (Illumina, software v1.5), according to manufacturer’s guidelines for scRNA-seq and scATAC-seq, respectively.

### scRNA-seq Data Preparation

The Cell Ranger pipeline was used for processing all samples post-sequencing (10X Genomics, version 3.1). Samples were demultiplexed and fastq files were generated using cellranger mkfastq. All samples were then individually aligned to the mouse reference genome mm10 using cellranger count with the intron = true flag to be consistent with single nuclei RNA-seq projects for which intronic reads are also kept. To account for varying sequencing read depth, we further ran cellranger aggr to normalize all samples to mapped read depth of the least sequenced sample, as previously described^4^. A single counts matrix for all samples is used as input for computational analysis using Seurat version 3.2.2^5^. Selected output quality control metrics from Cellranger count and Cellranger aggr for all samples are found in Table S1.

#### scRNA-seq Data Analysis

##### WT C57Bl6/J (E7.75 – E11.5) and Tbx1 WT and KO (E9.5 – E11.5)

Read-normalized aggregated values from cellranger aggr were used as initial input using the R package Seurat (v3.1) using the functions Read10X and CreateSeuratObject with standard variables. Each sample used in aggregation was identified by timepoint and replicate and assigned a unique name in metadata as “gem.group”. Quality control filtering included removal of outliers due to number of features/genes (nFeature_RNA > 2000 & nFeature_RNA < 7500), UMI counts (<80,000) and mitochrondial percentage (<15%). Cell cycle scores were added using the function CellCycleScoring. SCT normalization was then performed with regression based on cell cycle scores (vars.to.regress = c(“S.Score”, “G2M.Score”)). PCA analysis and batch correction using FastMNN was then performed using split.by = “gem.group”. Clustering was then run using the functions RunUMAP, FindNeighbors, and FindClusters and the output UMAP graphs were generated by DimPlot. Marker genes were identified by the function FindAllMarkers with standard settings. After initial processing, iterative rounds of filtering poor quality clusters and re-running clustering workflows. Cluster annotation, based on expression of known marker genes, was performed, leading to 11 broadly assigned cell types. Differential gene expression was performed using the Wilcoxon test between two groups with the function FindMarkers (logfc.threshold = 0.25 and min.pct = 0.1).

##### Venn Diagrams

All Venn diagrams were created with the R package VennDiagram using the function draw.pairwise.venn, manually settings the values for two areas and cross section^6^.

##### CellChat

Cell signaling analysis was performed using the R package CellChat^7^. Seurat objects containing specific cell types and time points of interest were imported to CellChat using the function GetAssayData(wt_cellchat, assay = “SCT”, slot = “data”) and createCellChat. All ligand-receptor and signaling pathways within the CellChatDB.mouse were kept for analysis. Initial preprocessing to identify over-expressed ligands and receptors was performed using the functions identifyOverExpressedGenes and identifyOverExpressedInteractions with standard settings. Inference of cell communication was calculated with computeCommunProb(cellchat_wt, raw.use = T, population.size = T) and filtered by filterCommunication(cellchat_wt, min.cells = 10). For downstream analysis, we identified all significant pathways for neural crest subsets as targets and all other cell types as senders using subsetCommunication. These results informed further subsetting for heatmaps and other visualizations. Pathway level cell communication was calculated with computeCommunProbPathway and aggregated networks were identified with aggregateNet, using standard settings. Network centrality scores were assigned with the function netAnalysis_computeCentrality. This workflow was run for both WT and KO datasets independently and differential signaling analysis was then run by merging the objects with mergeCellChat. A heatmap of differential signaling, based on significantly identified pathways from above, was generated using the function netAnalysis_signalingRole_heatmap and restricting the visualization to selected pathways. All scripts can be found on Github.

### scATAC-seq Data Preparation

#### Cell Ranger

The Cell Ranger ATAC workflow was used for all samples post sequencing (10X Genomics, version 1.2). Samples were first demultiplexed and fastq files were generated using the command cellranger-atac mkfastq. The command cellranger-atac count was then run to filter cell barcodes against a whitelist and align filtered fastq files to the mouse reference genome Mm10 using BWA-MEM. PCR duplicates and gel bead/cell multiplets were then removed (v1.2) and transposase cut sites were counted for each sample. Unlike scRNA-seq analysis, cellranger aggr was not run and instead the indexed fragments files (fragments.tsv.gz and fragments.tsv.gz.tbi) output from cellranger-atac count served as the inputs for computational analysis in ArchR^8^. Selected output quality control metrics from cellranger-atac count for all samples are found in Table S1.

#### scATAC-seq Data Analysis

##### WT C57Bl6/J (E7.75 – E11.5)

###### Project Generation in ArchR

Indexed fragments files for all samples were first used as inputs to generate a sample-specific Arrow file, including both sequencing-derived data and sample metadata. For each sample, the function createArrowFiles was run, removing cells with a TSS score below 6, and fragments below 1e4 and above 3e6. This step also generates a genome-wide TileMatrix using 500 bp bins and a GeneScoreMatrix, an estimated value of gene expression based on a weighted calculation of accessibility within a gene body and surrounding locus. Although Cell Ranger v1.2 implements cell multiplet removal, a second round of cell doublet removal was performed in ArchR using the functions addDoubletScores and filterDoublets. The number of cells counted as doublets and removed are found in Table S1. Each individual Arrow file (n=18 total) were then aggregated into a single ArchRProject for downstream analysis, leading to 64,956 cells, with a median TSS of 12.193 and median value of 48,812 fragments per cell.

##### Tbx1 WT and KO (E9.5 – E11.5)

Similar functions were run with the following exception: createArrowFiles, fragments between 1e4 and 5e6. After doublet removal, each individual Arrow file (n=13, 6 WT and 7 KO) was aggregated and the final project contained 78,062 cells with a median TSS 12.2 and median fragments of 47,353.

##### Tbx1 WT and KO (E8.5)

Similar functions were run with the following exception: createArrowFiles, fragments between 1e4 and 8e5. After doublet removal, each individual Arrow file (n=13, 4 WT and 3 KO) was aggregated and the final project contained 15,178 cells with a median TSS 11.38 and median fragments of 51,252.

#### Clustering

##### WT C57Bl6/J (E7.75 – E11.5)

After generation of an aggregated Project, dimensionality reduction was performed using ArchR’s implementation of Iterative Latent Sematic Indexing (LSI) with the function addIterativeLSI and the LSI method of “log(tf-idf)”. The 500 bp TileMatrix was used as input and 4 iterations of LSI were performed with 25,000 variable features, increasing resolution values (0.1, 0.2, and 0.4), and reduced dimensions 2:25. The reduced dimension values from IterativeLSI were then used as input for batch correction using Harmony with the function addHarmony. Clustering was then performed using Harmony corrected values with addClusters, invoking the function FindClusters, with a resolution of 0.8 from the R package Seurat. Finally, clusters were visualized with function plotEmbedding, using batch corrected single-cell embedding values from Uniform Manifold Approximation and Projection (UMAP) using the function addUMAP.

##### Tbx1 WT and KO (E9.5 – E11.5)

Iterative-LSI was performed with 3 iterations (resolutions 0.1 and 0.2) with reduced dimensions 2:30. Final clustering was performed with maximum clusters of 40 and resolution of 0.6.

##### Tbx1 WT and KO (E8.5)

Iterative-LSI was performed with 2 iterations (resolution 0.2) and reduced dimensions 2:30. Final clustering included resolution value of 0.6

#### Cluster Annotation

Determination of the identity of each cluster in scATAC-seq was performed manually after consideration of combined information from GeneScore plots and scRNA-seq integration. A list of known marker genes for each broadly annotated cluster from scRNA-seq analysis were used to generate GeneScore plots in the scATAC-seq data with the function plotEmbedding. The R package MAGIC was used to impute GeneScore values for each gene (PMID 29961576). For scRNA-seq integration, Canonical Correlation Analysis using Seurat’s FindTransferAnchors function was performed by addGeneIntegrationMatrix. Manual inspection of GeneScore plots and scRNA-seq integration, along with iterative rounds of re-clustering, was then performed along and clusters were annotated individually using the revalue function from the R package plyr.

A Jaccard Similarity Analysis using the predicted integration label of clusters from scRNA-seq and scATAC-seq integration and the manual annotation was run similarly to previous methods^9^. Briefly, a similarity index was calculated by the proportion of cells with similar labels between both assignments over the number of cells with either of the two labels. We then visualized the result proportions with the R package ComplexHeatmap function pheatmap^10^. Similar methods were used for Tbx1 analysis.

#### Peak Calling and Motif Enrichment

After cluster annotation, pseudo-bulk replicates of cells within similar groups were created to facilitate peak calling. Replicates were created using addGroupCoverages, using 2-18 replicates per annotated cluster, max fragments of 7.5e7, a sample ratio of 0.8 and filtering for a minimum and maximum of 20 and 3000 cells, respectively. Peak calling was then performed using addReproduciblePeakSet with a maximum of 2500 peaks per cell and 2e5 peaks per cluster. This function invokes MACS2 using the following criteria: “--shift = −75, --extsize = 150, --nomodel, --keep-dup all, --q 0.1, --nolambda” and excluding mitochondrial chromosome genes and chromosome Y^11^. We then used ArchR’s iterative overlap peak merging method to create a union peakset of 548,312 unique peaks.

#### Tbx1 WT and KO (E8.5)

Similar methods were used with exceptions: addReproduciblePeakSet with 2000 peaks per cell and a max of 150,000 peaks. A final union peak set of 481,439 peaks were generated.

Cluster-enriched marker peaks were then identified with getMarkerFeatures, using a wilcoxon test and normalizing for biases from TSS enrichment scores and sequencing depth, and visualized with plotMarkerHeatmap, filtering for FDR <= 0.05 and Log2 Fold Change of >= 1.5. Motif enrichment of cluster enriched peaks was done using addMotifAnnotations with the Homer database of motifs^12^. Enriched motifs per cluster were visualized by first running peakAnnoEnrichment, with FDR <= 0.05 and Log2FC >=1 and then subsetting for manually selected, non-redundant motifs for each cluster and finally creating a heatmap with the draw function in ComplexHeatmap. Motif enrichment at single cell resolution was done using the R package ChromVAR by first adding background peaks (addBgdPeaks) and then computing z-scored deviations of each motif per cell with addDeviationsMatrix^13^. ChromVAR motif enrichments per cell were visualized with batch corrected UMAP embeddings in plotEmbedding.

#### Peak -to-Gene Linkage

Peak-to-gene linkage analysis was performed in ArchR using the addPeak2GeneLinks command, using the batch corrected Harmony embedding values and removing cells with predictionCutOff < 0.4. This filtering only discarded 421 out of 64,956 total cells, highlighting the strong concordance of cluster annotation for cells in scATAC-seq and scRNA-seq. A total of 67,606 linkages were found using FDR < 1e-04, corCutOff = 0.5 and a resolution of 1. These peaks associated with each linkage were then exported as a bed file and inputted to homer for annotation to nearest gene.

#### Peak Annotation

The R package ChIPpeakAnno was used to determine the genomic annotation of each peak in the dataset and the distance to known TSS^14^. We first converted the 548,312 total unique peaks to a GRanges object using the function toGRanges and then performed peak annotation using the command annotatePeak, setting tssRegion to 2kb surrounding the TSS and using the Mm10 reference genome. Distances to known TSS was calculated using the plotDistToTSS command.

#### HOMER

Motif enrichment was performed using the function findMotifsGenome.pl -size given -mask. Annotation to nearest gene was performed using the function annotatePeaks.pl.

#### Subsetting and designation of genotype

After cluster annotation, specific clusters for any given dataset were chosen for refined subsetted analysis by using the R package plyr and base R functions. First samples or clusters were marked with a unique string by the function substr. These vector values were then assigned as a metadata column to each cell using the functions as.character(vector.of.cells). For subsetting specific clusters, similar functions were used and clusters to keep were given a value of “True” and then filtered using BioGenerics::which (object %in% “True”).

#### Statistical Test for Comparing Cell Proportion Differences

A statistical test comparing whether loss or gain of proportions of cells for a given cluster between WT and KO was performed using scProportionTest: https://github.com/rpolicastro/scProportionTest. Briefly, a Monte-carlo/permutation test was run by pooling cells for each cluster together and randomly assigning cells back to a given condition and re-calculating proportions. The identities for cell type were previously defined in Seurat and this randomized process was performed 10,000 times. Differences in magnitude greater than chance and confidence intervals are measured for each cell type, comparing genotypes (FDR < 0.05, abs(Log2FD > 0.58)).

### Machine Learning Algorithm for Enhancer Predictions

Logistic regression classifiers (L1 penalty, LASSO) were trained on validated mouse enhancers from the VISTA Enhancer Browser. A cardiac-specific model used validated heart enhancers as positives, and non-heart enhancers as well as candidates that failed to validate as negatives. A second neural crest-specific model used validated branchial arch and facial mesenchyme enhancers as neural crest positives and non-neural crest tissue enhancers as negatives. All enhancers were featurized using intersections (bedtools) with publicly available data including ENCODE mouse E10.5 reference epigenomes, Akerberg et al. 2019 E12.5 cardiomyocyte ATAC- and ChIP-seq^15^, Cesario et al. 2015 E11.5 maxillary arch ChIP-seq^16^, Elliott et al. 2020 E11.5 mandibular prominence and whole face ChIP-seq^17^, Haro et al. 2017 E12.5 limb ChIP-seq^18^, Infante et al. 2013 E11.5 hindlimb ChIP-seq^19^, Lex et al. 2020 E10.5 forelimb ChIP-seq,^20^ Luna-Zurita et al. 2016 mES cardiomyocyte ChIP-exo^21^, and Zhou et al. 2017 E12.5 forebrain and heart ChIP-seq^22^, as well as average sequence conservation over the region (phyloP/PhastCons 100-way)^23^. The trained classifiers scored all scATAC-seq peaks between 0 and 1, estimating their similarity to either validated VISTA heart enhancers (model 1) or neural crest enhancers (model 2). A decision threshold (0.4 for both models) was then applied to turn this continuous score into a binary prediction (‘positive’ or ‘negative’).

Non-ENCODE data was reprocessed using ENCODE pipelines for ATAC-seq, DNase-seq, and ChIP-seq datasets, while ChIP-exo data was reprocessed using GEM. All data was aligned to mm10.

### Zebrafish Transgenic Reporter Assay

Putative enhancer regions were amplified from genomic DNA using primers listed below and were cloned into the E1b-Tol2-eGFP vector. 25 ng E1b-Tol2-eGFP plasmid carrying non-coding fragment and 125ng Tol2 mRNA were injected into wildtype, cmlc2:mCherry and sox10:mRFP eggs at the one-cell stage as previously described^24^. F0 founder embryos were raised to 72 hpf. Cloning Primers:

1. chr16:96,212,776-96,213,475 CAGGCCAGATGGGCCCtcgaGtcatgcaggaagagggaactt CCCTCTAGAGTCGAGAgatcTTCAACTCATCATCGATAGACAAGT
2. chr3:152,133,489-152,134,198 CAGGCCAGATGGGCCCtcgaGCTATCTGCATTTTAAAGTAGACTGTGT CCCTCTAGAGTCGAGAgatcTCTGCAGTCTCTGCTTTTCTCGGT
3. chr4:135,313,196-135,313,841 CAGGCCAGATGGGCCCtcgaGGCCATCTCTAGCTTTCAGTTCA CCCTCTAGAGTCGAGAgatcTTGCTATGACTACAGGGATAAGGGGT
4. chr1:77,338,250-77,338,750: CAGGCCAGATGGGCCCtcgaGCGTGACTCTTGAGGCAGATGCT CCCTCTAGAGTCGAGAgatcTCTTCTAATGAGGGTTGGCAGGATATCT
5. chr16:30,627,503-30,628,003 CAGGCCAGATGGGCCCtcgaGCTAAGCTGTAGGAATACATTGTTGTT CCCTCTAGAGTCGAGAgatcTCTAAGATAGTATAAGGATCTAGACTTG
6. chr14:100,735,266-100,735,766 CAGGCCAGATGGGCCCtcgaGGCCATTTTCTGACTTTCTTCTATCTC CCCTCTAGAGTCGAGAgatcTGGATGCATTTTGTCTAACATCTTTCA

#### Imaging

In glass-bottom Petri dishes, live embryos were mounted for confocal imaging using 0.5-1% low-melting agarose (Sigma, cat. no. A9414). Fixed embryos for confocal imaging were de-yolked using lash tools and mounted glass slides with coverslips. All confocal imaging was performed on a Zeiss 517 LSM 710 FCS confocal microscope.

### Embryo Dissection for RNAscope Assay

Embryos meant for RNAscope assay were collected through timed mating similar to methods for single cell assays. After dissections, embryos were fixed overnight with 4% paraformaldehyde (50-980-487, Fisher Scientific). The next day, embryos were washed in PBS and dehydrated in 70% EtOH overnight. Embryos were then embedded in Histogel to maintain correct orientation prior to parafin processing (22-110-678, Fisher Scientific). Embryos were processed in paraffin overnight and sectioned onto glass slides at 5 um thickness.

### RNAscope In Situ Hybridization

Fluorescence in situ hybridization was performed using RNAscope assay (ACDbio) according to manufacturer’s protocols. Probes used were: Hand1 (429658-C2), Smoc1 (571511-C2), Hoxb3 (51585) and Zsgreen (461258). All slides were imaged on Olympus FV3000RS confocal microscope.

### Luciferase Assay

Fasta sequences of candidate enhancers were obtained by inputting mm10 coordinates in command line using twoBitToFa command. Candidate enhancers were PCR amplified (Takara PrimeSTAR GXL premix, R051A) from mouse genomic DNA using primers listed below. Following agarose gel purification (NucleoSpin Gel and PCR clean up mini kit, 740609), PCR products were cloned into pGL4.23 vector (Promega, E841A) digested with XhoI and HindIII (New England Biolabs, R0146S and R3104S, respectively) using Cold Fusion cloning (System Biosciences, MC010B-1). All constructs were sequence verified for successful insert integration, scaled up and purified prior to transfection (Qiagen HiSpeed Maxi prep, 12663). For luciferase assays (Promega, Dual-Luciferase Reporter Assay System, E1960), HeLa cells (ATCC, CCL-2) were seeded at 70,000 cells per well of a 24 well plate 24h prior to transfection. All enhancer plasmids (200 ng) for testing were co-transfected with Renilla control plasmid (20 ng) and either a plasmid containing mouse Tbx1 cDNA (200 ng, Origene, PS100001) or empty vector control (200ng) with 2.4 ul Fugene HD (Promega, E2311) in 43 ul Opti-MEM media (Gibco, 31985-062). Each construct was transfected in quadruplicate technical replicates and at least 3 biological replicates (separate experiments) were performed. Luciferase assays were performed 24-36 hours after transfection according to manufacturer’s protocols. Measurements were taken using SpectraMax MiniMax 300 imaging cytometer with SoftMax Pro6, Version 6.4 (Molecular Devices).

Sema3c, Peak 1, chr5: 17551355 – 17552016
CCTGAGCTCGCTAGCCTCGAGAACTTAGAGTCATGCCTTCCATTCT
TACCCTCTAGTGTCTAAGCTTACAGGGCAAAGCCAGGGAGAA

Sema3c, Peak 2, chr5: 17744105 – 17744813
CCTGAGCTCGCTAGCCTCGAGATGTGGATCCTTGATCTGTAGT
TACCCTCTAGTGTCTAAGCTTGAAATTTGAACAATTCTCTTATGA

### Data Availability and Scripts

All scripts used for analysis have been deposited to GitHub: https://github.com/sanjeevRJMU1. Also included on GitHub are all peak calls and enhancer predictions (bed files). A publicly accessible UCSC genome browser session for WT scATAC-seq peaks and enhancer predictions can be found here: https://genome.ucsc.edu/s/sanjeevranade%40gmail.com/SSR_SrivastavaLab_scATAC. Raw sequencing data and processed data for scRNA-seq and scATAC-seq for WT and WT/Tbx1 combined analysis are publicly available through NCBI GEO: https://www.ncbi.nlm.nih.gov/geo/query/acc.cgi?acc=GSE198567

## Acknowledgements

We would like to thank the center for advanced technology (CAT) for sequencing; the Gladstone Histology and Light Microscopy Core; and the Gladstone Stem Cell Core. We acknowledge Francoise Chanut for her editorial input and Giovanni Maki for graphical illustrations (Gladstone Institutes), and Bethany Taylor for assistance in preparing the manuscript. D.S. is supported by NIH/NHLBI P01 HL098707, P01 HL146366, R01 HL057181, R01 HL127240, and by the Roddenberry Foundation, the L.K. Whittier Foundation, and the Younger Family Fund. S.R. is a Winslow Fellow funded by contributions from Clark and Sharon Winslow. T.N. is supported by the Japan Society for the Promotion of Science Overseas Research Fellowship. A.P. is supported by NIH K08 HL157700, Sarnoff Cardiovascular Research Foundation and Michael Antonov Charitable Foundation. K.S.P. is supported by NIH P01 HL098707, P01 HL146366, UM1 HL098179, and the San Simeon Fund.

## Author Contributions

S.S.R. and D.S. designed and directed all experiments in this study. S.W. and K.S.P. designed and generated machine learning models for enhancer predictions. I.Z. and B.L.B. designed and performed zebrafish transgenic enhancer reporter experiments. S.S.R., T.N., B.v.S., L.Y., L.G.W., Y.H. performed all mouse embryonic experiments for scATAC-seq and scRNA-seq. B.v.S. and S.S.R. performed embryonic staining experiments and microscopy. S.S.R. performed all computational analysis of single cell experiments with input and direction from A.P., P.P., M.A., M.W.C. and C.A.G. Luciferase assay experimented were performed by L.Y., S.S.R., and B.G.T. S.S.R. and L.Y. designed and cloned all transgenic reporter constructs. S.S.R. and D.S. wrote the manuscript.

## Additional Information

Three supplementary tables are provided.

Corresponding author: Deepak Srivastava

