## Supplemental Figures for "Single Cell Epigenetics Reveal Cell-Cell Communication Networks in Normal and Abnormal Cardiac Morphogenesis"

### Extended Data Figures and Legends

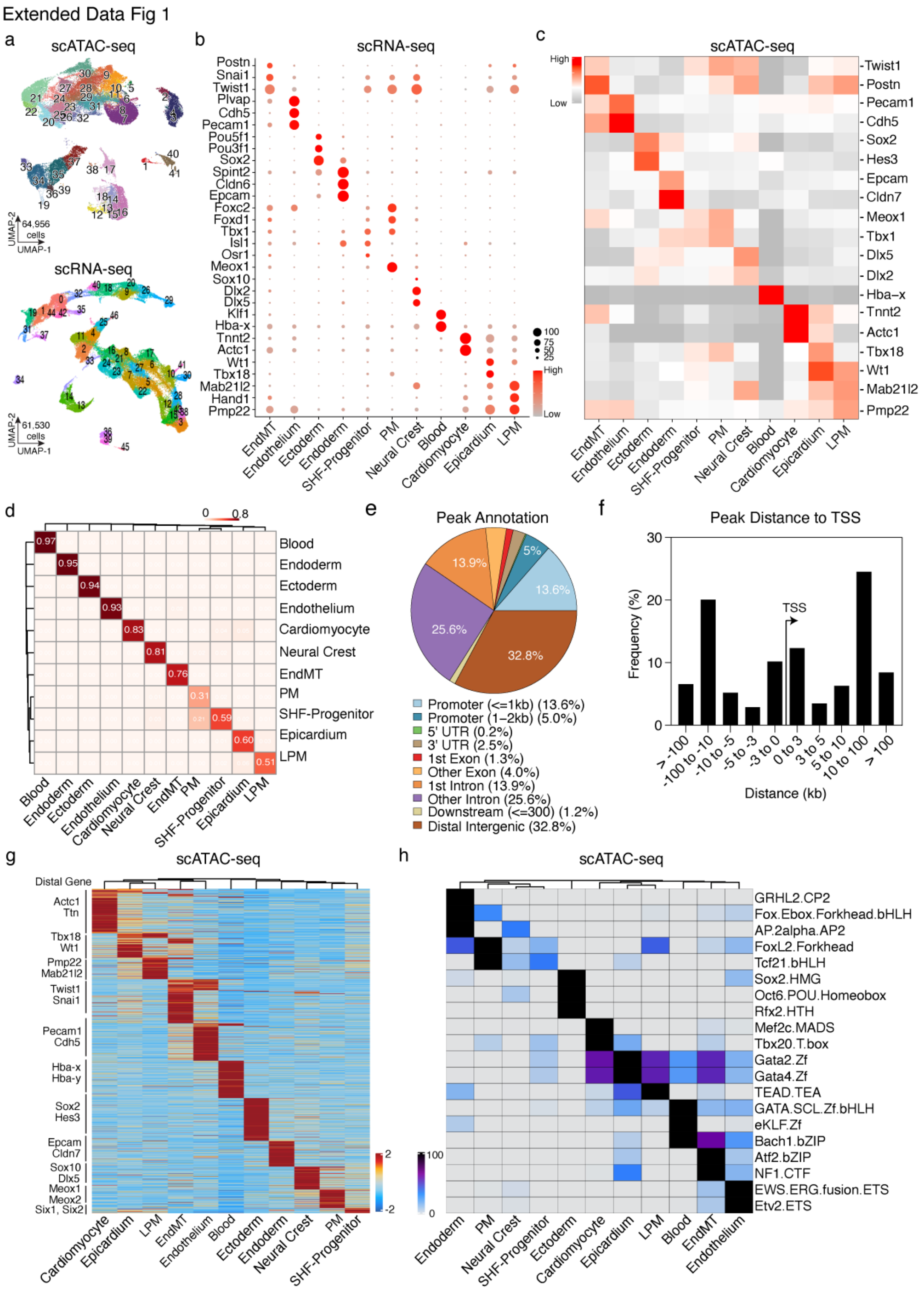


**Extended Data Fig. 1 | Integrated Single Cell Multimodal Analysis of Heart Development.** **a,** UMAP of scATAC-seq and scRNA-seq of cardiac and pharyngeal arch development from E7.75 to E11.5 colored by clusters. scATAC-seq UMAP contains 41 clusters with 64,956 cells (n=18 embryos) and scRNA-seq contains 46 clusters and 61,530 cells (n=17 embryos). **b,** Dot plot of gene expression for curated list of marker genes (scRNA-seq) enriched for each cluster. Dot size represents proportion of gene expression within cells of that cluster and dot color represents magnitude of expression (grey to red, low to high levels). EndMT, Endothelial to Mesenchymal Transformation; PM, paraxial mesoderm; SHF Progenitor, second heart field progenitor; LPM, lateral plate mesoderm. **c,** Heatmap of gene score values for curated set of marker genes applied from previous knowledge and scRNA-seq outputs. Values for each cluster represent z-scores ranging from -1 to 2 (low to high scale colored by grey to red). **d,** Heatmap of Jaccard index similarity analysis post Seurat v3 canonical correlation analysis transfer labelling. Values range from 0.1 – 1.0 and color corresponds to concordance of label matching between modalities. **e,** ChipPeakAno-based annotation of scATAC-seq peaks based on genomic location. **f,** Bar plot of peaks binned by distance from known transcription start sites. **g,** Heatmap of cluster-enriched peaks (FDR <= 0.5 & Log2FC >= 1.5). Peaks were assigned to nearest gene (HOMER) and curated set of known marker genes are labelled on the side. Colors represent z-scored values for enrichment of peak to a cluster. **h,** Heatmap of motifs enriched within each cluster using ArchR's peakAnnoEnrichment function with the HOMER database (FDR <= 0.5 & Log2FC >= 1). Color of cells scaled by normalized enrichment (-log10(p-adj)) ranging from 0-100.

Extended Data Figure 2

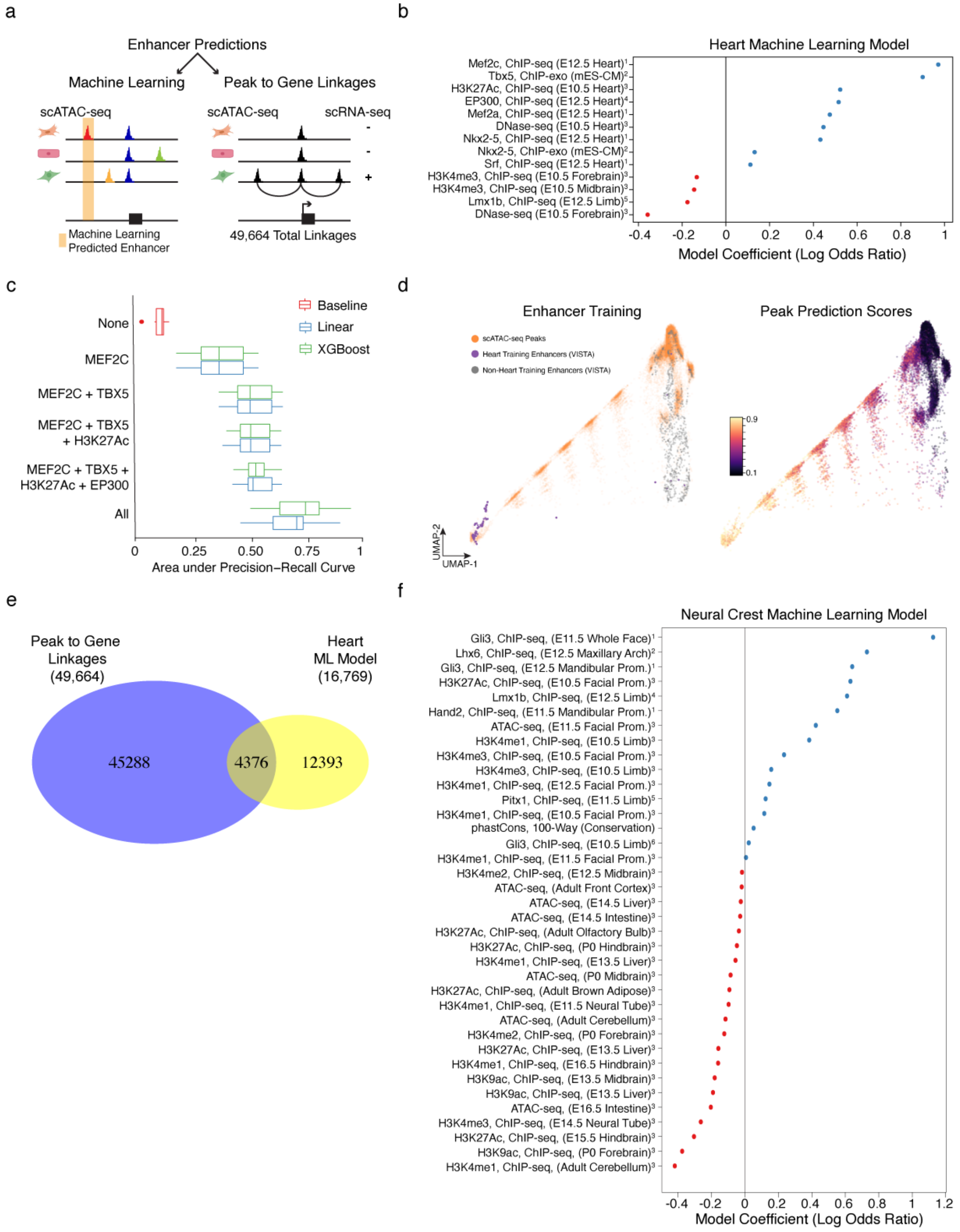


**Extended Data Fig. 2 | Characteristics Underlying Machine Learning Models. a,** Cartoon illustration of two methods for enhancer predictions from scATAC-seq data. One method utilizes the integrated multimodal matrix to assign putative chromatin accessibility peaks to genes using

the ArchR peak to gene workflow (corCutOff=0.5, FDRcutoff = 1e-04, varCutOffATAC and varCutOffRNA = 0.25, resolution = 1). The machine learning method is an orthogonal approach for assigning values to all peaks for predicted enhancer activity. **b,** For the heart machine learning model, 13 feature inputs retrained post model training and respective model coefficients (log odds ratio). References for each dataset used are described in methods section.
**c,** Benchmark comparison of performance for heart model using luck (baseline), a penalized linear model or an alternative XGBoost method. Model performance, assessed by area under the precision-recall curve, was tested by selecting individual high performing datasets or in combinations. **d,** UMAP representations depicting clear separation of cardiac model performance in assigning mouse heart scATAC-seq peaks compared to known VISTA heart and non-heart enhancers. Peak prediction scores with highest values (from 0-1) were clustered closely with heart enhancers and show a gradient of values correlating with proximity to heart enhancers. Each dot in UMAPs is a genomic locus (peak). **e,** For the neural crest machine learning model, 37 feature inputs retrained post model training and respective model coefficients (log odds ratio). References for each dataset used are described in methods section.

Extended Data Figure 3

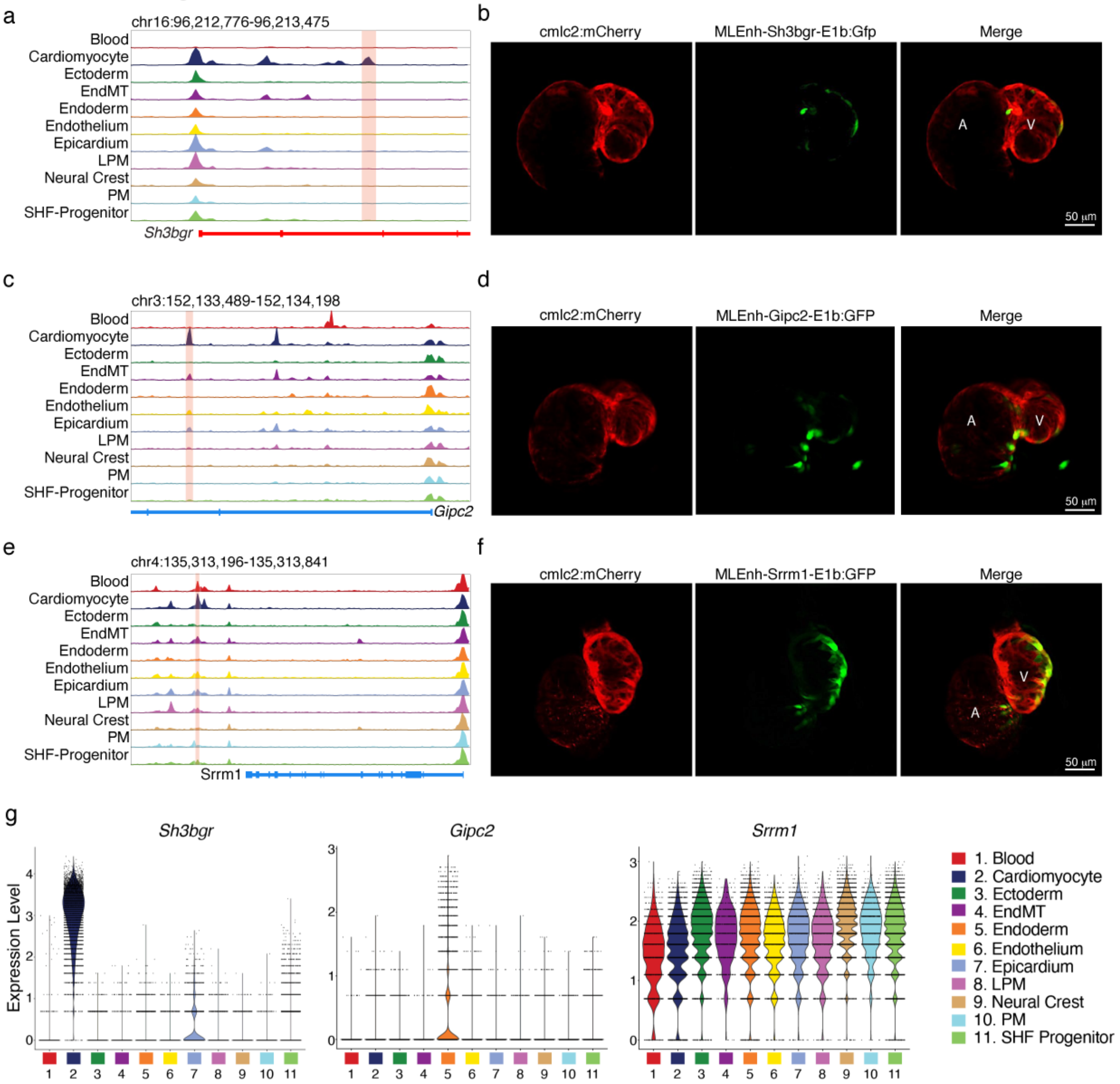


**Extended Data Fig. 3 | Experimental Validation of Select Predicted Cardiomyocyte Enhancers. a,** Browser track view of genomic locus surrounding Sh3bgr (mm10:chr16:96200470-96228933) and scATAC-seq signal per cluster. Highlighted region (chr16:96,212,776-96,213,475) is a highly predicted cardiomyocyte enhancer. **b,** Epifluorescence imaging of transgenic cmlc2:mCherry F0 Zebrafish heart at 72hpf injected with predicted Sh3bgr intronic enhancer upstream of a minimal E1b-promoter driven GFP. Total 16/42 embryos GFP positive. Scale, 50 µm. **c,** Browser track view of genomic locus surrounding Gipc2 (chr3:152093841-152165900) and scATAC-seq signal per cluster. Highlighted region (chr3:152125899−152166900) is a highly predicted cardiomyocyte enhancer. **d,** Epifluorescence imaging of transgenic cmlc2:mCherry F0 Zebrafish heart at 72hpf injected with predicted Gipc2 intronic enhancer upstream of a minimal E1b-promoter driven GFP. Total 16/42 embryos GFP positive. Scale, 50 µm. **e,** Browser track view of genomic locus surrounding Srrm1 (chr4:135320484-135353214) and scATAC-seq signal per cluster. Highlighted region (chr4:135303213−135354214) is a highly predicted cardiomyocyte enhancer. **f,** Epifluorescence imaging of transgenic cmlc2:mCherry F0 Zebrafish heart at 72hpf injected with predicted Srrm1 distal enhancer upstream of a minimal E1b-promoter driven GFP. Total 11/33 embryos GFP positive. Scale, 50 µm. **g,** Violin plots of Sh3bgr, Gipc2 and Srrm1 in all populations.

Extended Data Figure 4

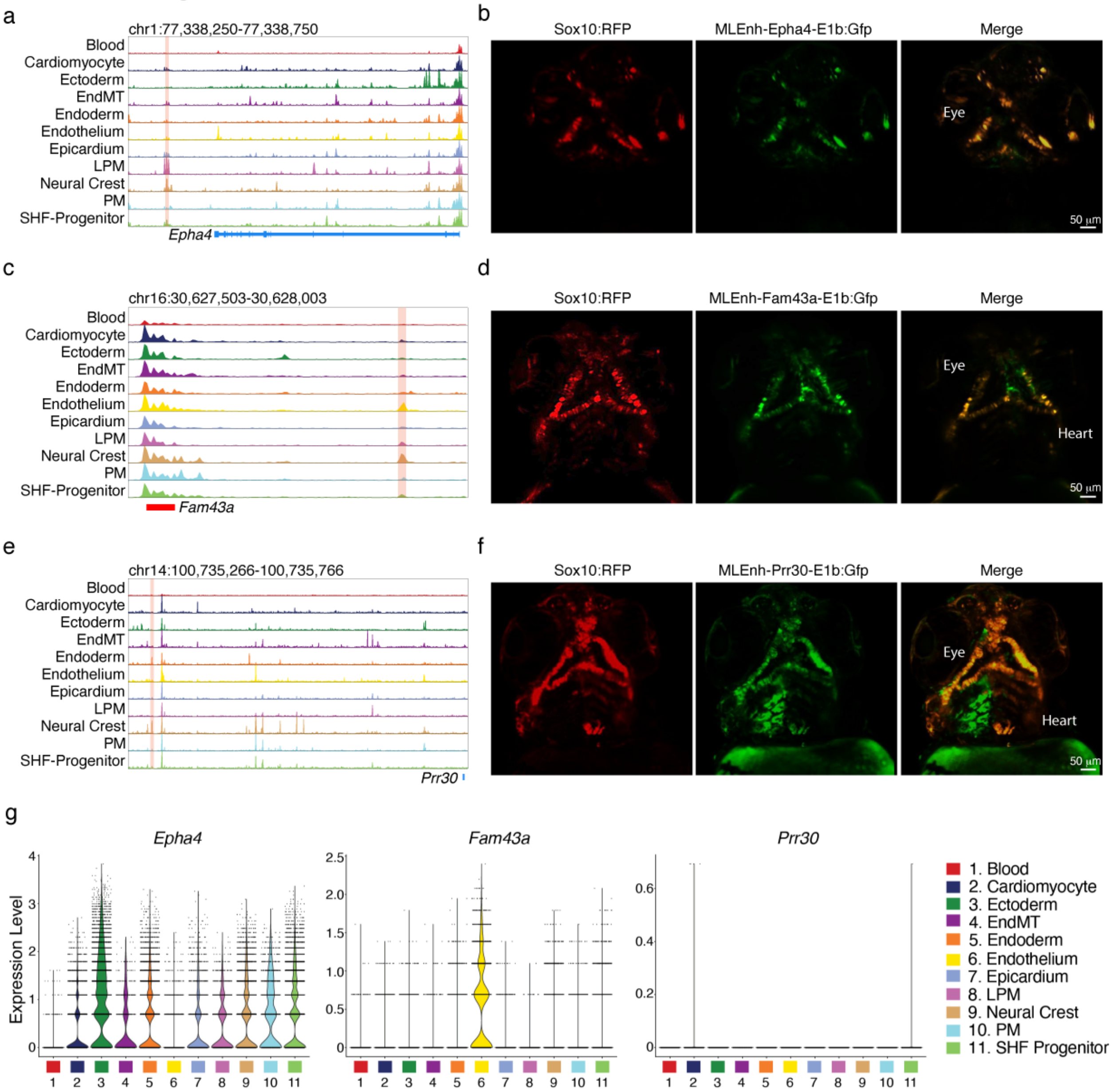


**Extended Data Figure 4 | Experimental Validation of Select Predicted Neural Crest Enhancers. a,** Browser track view of genomic locus surrounding Epha4 (mm10:chr1:77,367,185-77,515,088) and scATAC-seq signal per cluster. Highlighted region (chr1:77,338,250-77,338,750) is a highly predicted neural crest enhancer. **b,** Epifluorescence imaging of transgenic sox10:mRFP F0 Zebrafish heart at 72hpf injected with predicted Epha4 distal enhancer upstream of a minimal E1b-promoter driven GFP. Total 8/21 embryos GFP positive. Scale, 50 µm. **c,** Browser track view of genomic locus surrounding Fam43a (chr16:30,599,723-30,602,797) and scATAC-seq signal per cluster. Highlighted region (chr16:30,627,503-30,628,003) is a highly predicted neural crest enhancer. **d,** Epifluorescence imaging of transgenic sox10:mRFP F0 Zebrafish heart at 72hpf injected with predicted Fam43a distal enhancer upstream of a minimal E1b-promoter driven GFP. Total 2/5 embryos GFP positive. Scale, 50 μm. **e,** Browser track view of genomic locus surrounding Prr30 (mm10: chr14:101,197,690-101,200,069) and scATAC-seq signal per cluster. Highlighted region (chr14:100,735,266-100,735,766) is a highly predicted neural crest enhancer. **f,** Epifluorescence imaging of transgenic sox10:mRFP F0 Zebrafish heart at 72hpf injected with predicted Prr30 distal enhancer upstream of a minimal E1b-promoter driven GFP. Total 11/33 embryos GFP positive. Scale, 50 μm. **g,** Violin plots of Epha4, Fam43a and Prr30 in all populations.

Extended Data Figure 5

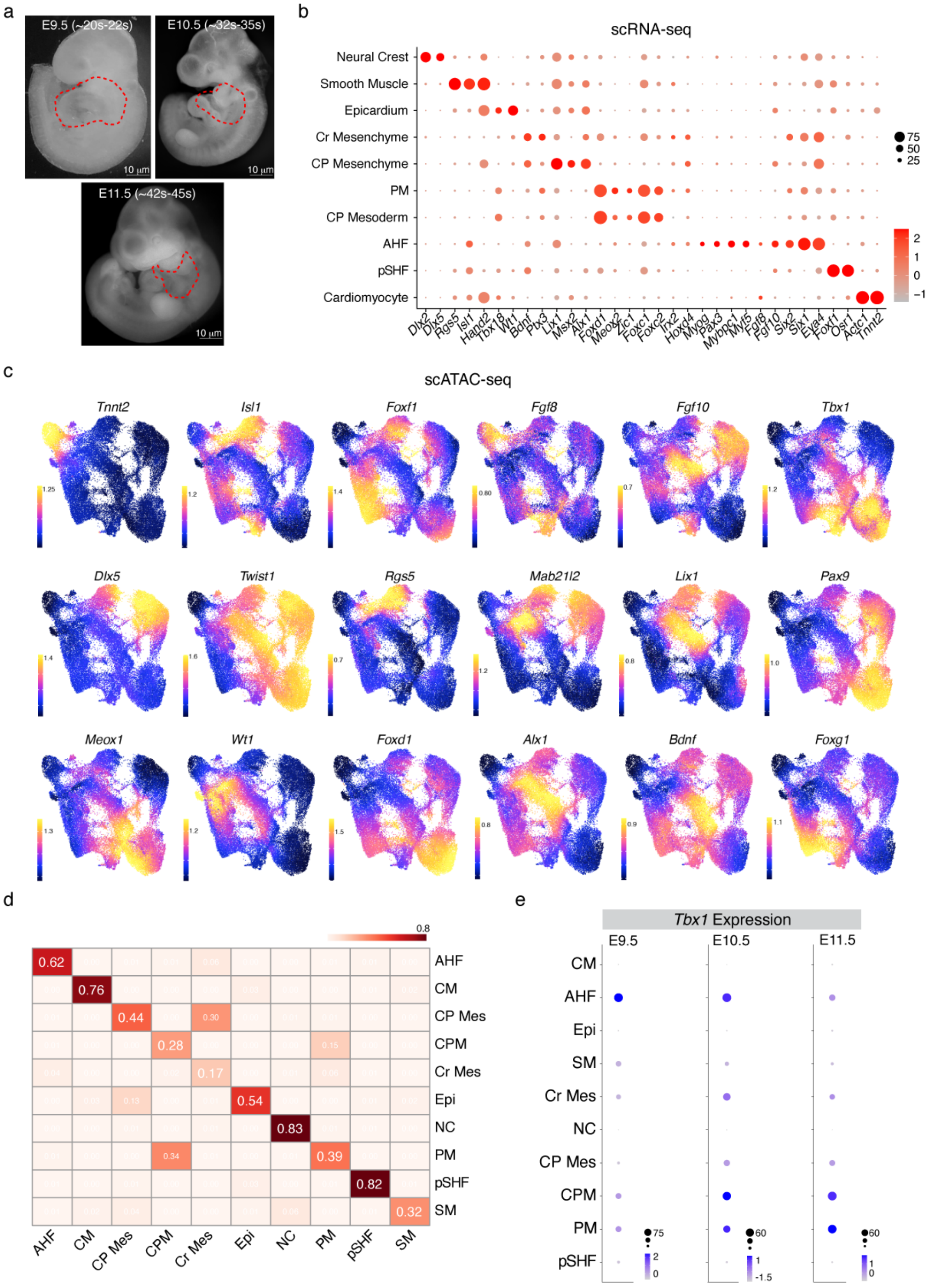


**Extended Data Fig. 5 | Integrated scRNA-seq and scATAC-seq of Tbx1 WT and KO Embryos at E9.5 – E11.5. a,** Representative images of embryos at three stages of analysis: E9.5, E10.5 and E11.5. Dotted lines are the regions micro-dissected for both modalities. **b,** Dot plot of gene expression values (scRNA-seq) for marker genes of mesoderm and neural crest lineages.

Color intensity represents scaled expression levels and dot size is proportion of cells within cluster expressing the gene. **c,** UMAP images of gene score values (scATAC-seq), imputed values of chromatin accessibility within gene bodies and surrounding locus. Colors represent log2 normalized values + 1. **d,** Jaccard index heatmap of cell type annotation comparison of scATAC-seq and scRNA-seq. Scaled values (scale of 0-1) are represented. **e,** Dot plot of Tbx1 expression (scRNA-seq) in all cell types at three stages (E.95, left; E10.5, center; E11.5, right). Color intensity represents scaled expression levels and dot size is proportion of cells within cluster expressing the gene.

Extended Data Figure 6

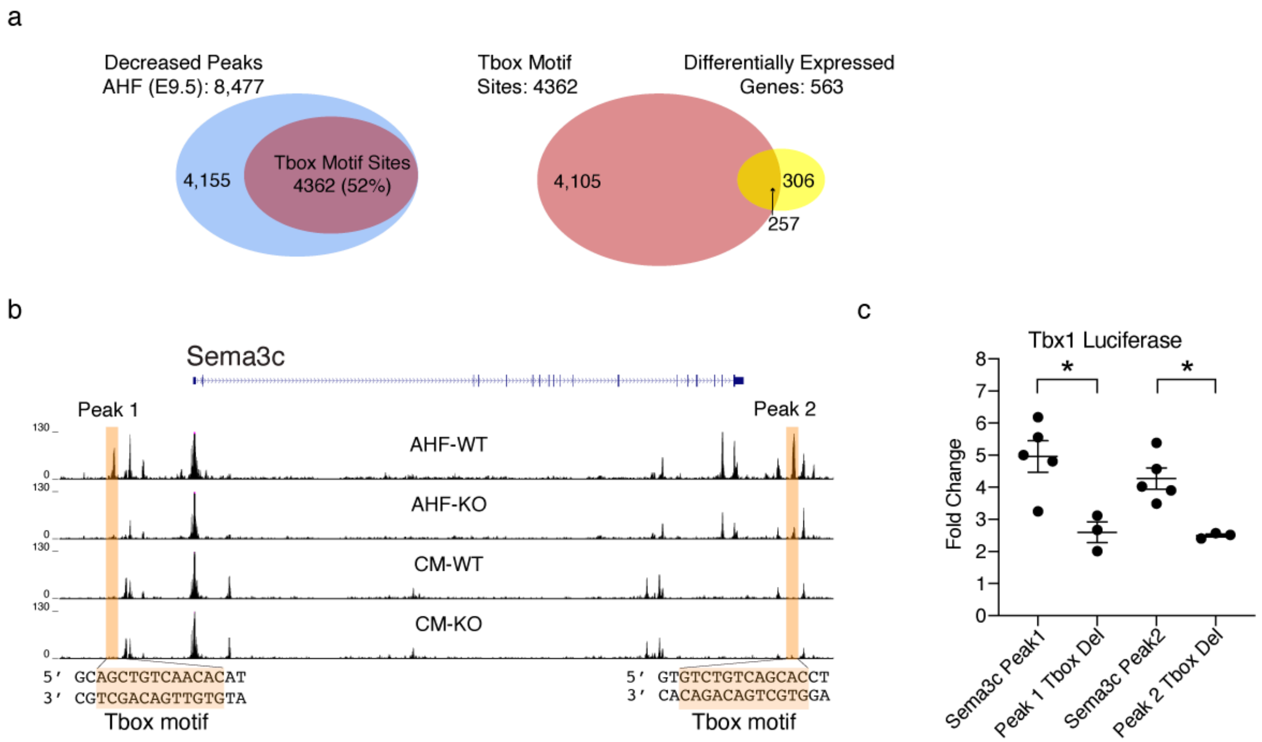


**Extended Data Fig. 6│ Intersection and Experimental Validation of Differentially Accessible Sites and Expressed Genes. a,** Venn diagram of differentially accessible sites lost in AHF cells at E9.5 containing a putative Tbox motif (scATAC-seq, HOMER) and differentially expressed genes (scRNA-seq). **b,** Browser track view of Sema3c locus with highlighted regions representing two distal chromatin accessibility peaks lost selectively in AHF cells at E9.5 in

Tbx1 KO embryos. 12 bp Tbox motif sequence in reverse complement orientation (3' to 5'), as indicated by HOMER analysis, underlined and shown below peaks. **c,** Luciferase assay results of both Tbx1-dependent peaks distal to Sema3c in the presence of a cDNA expression vector containing mouse Tbx1 compared to peaks with a 12 bp deletion of the putative Tbox motif. Data shown with WT sequence normalized to 1, n=3, * $p < 0.5$. Error bars indicate S.E.M.

Extended Data Figure 7

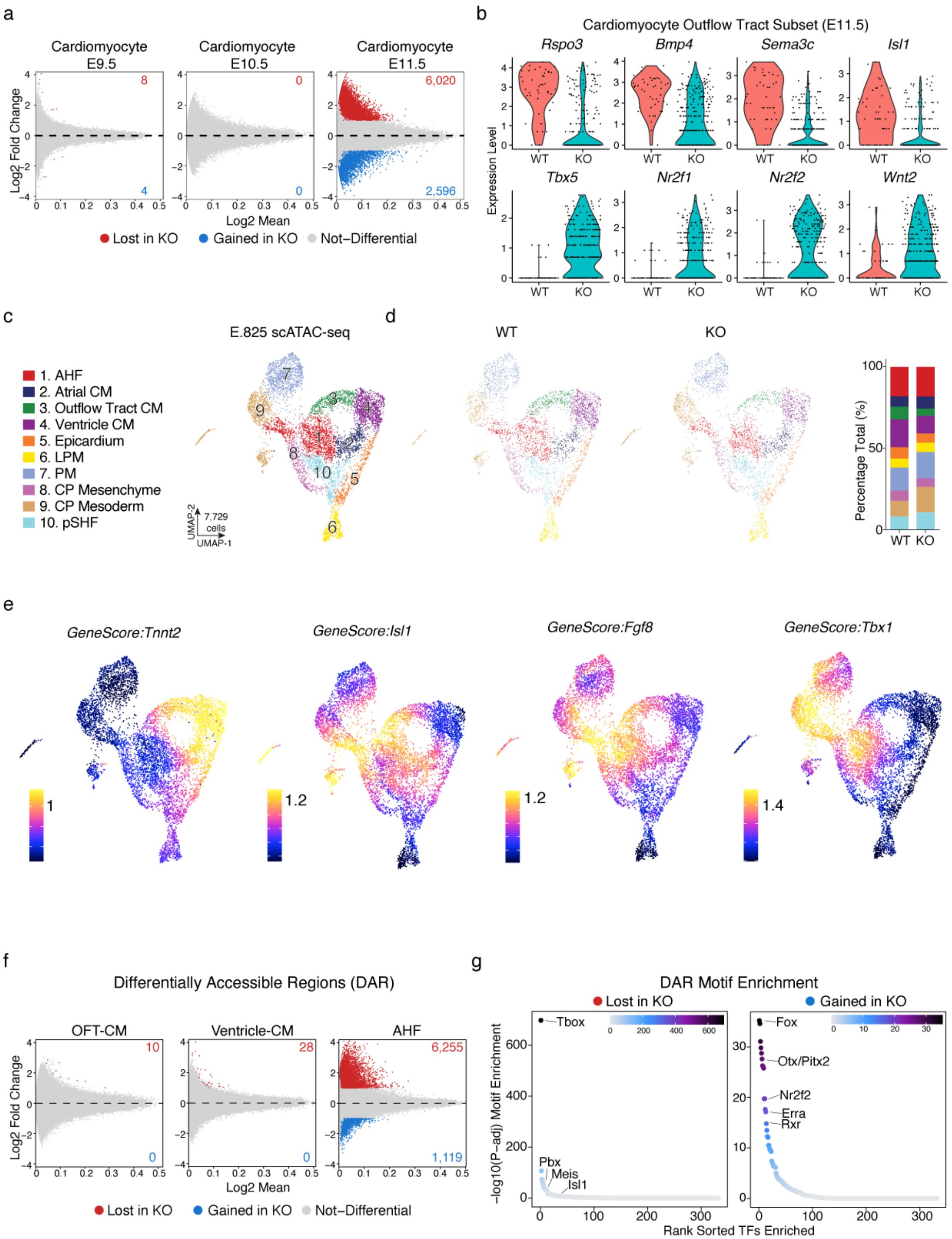


**Extended Data Fig. 7│ Tbx1 Loss is Dispensable for Early Cardiomyocyte Specification But Alters Outflow Tract Identity at Mid-gestation. a,** PMA plots of differential accessibility comparing Tbx1 WT and KO within cardiomyocyte population at E9.5, E10.5 and E11.5 (log2

Fold change over log2 mean intensity, FDR <= 0.05 & abs(Log2FC) >= 1). Number of DARs identified plotted within volcano plot. **b,** Violin plots of log normalized values for E11.5 cardiomyocytes, subsetted to outflow tract myocardial cells, comparing WT and KO for outflow tract and atrial myocardial genes. **c,** Annotated UMAP of scATAC-seq cells at E8.5, subsetted to contain only mesoderm lineage populations (n = 7,729 cells) including both WT and KO cells (n= 4 WT and 3 KO reps). AHF, anterior heart field; CM, cardiomyocyte; LPM, lateral plate mesoderm; PM, paraxial mesoderm; CP, cardiopharyngeal; pSHF, posterior second heart field. **d,** UMAP images and bar plot of samples for WT (4,080 cells) and KO (3,649 cells). **e,** UMAP images of gene score values (scATAC-seq), imputed values of chromatin accessibility within gene bodies and surrounding locus. Colors represent log2 normalized values + 1. **f,** PMA plots of differential accessibility comparing Tbx1 WT and KO within AHF, outflow tract and ventricular myocardium at E8.5 (log2 Fold change over log2 mean intensity, FDR <= 0.05 & abs(Log2FC) >= 1). Number of DARs identified plotted within volcano plot. **g,** Plot of motifs enriched within sites lost and gained in KO in AHF at E8.5. Rank sorted TF motifs on x-axis and -log10 FDR of motif enrichment on y-axis. Scale bar represents mlog10 adjusted p-value range.

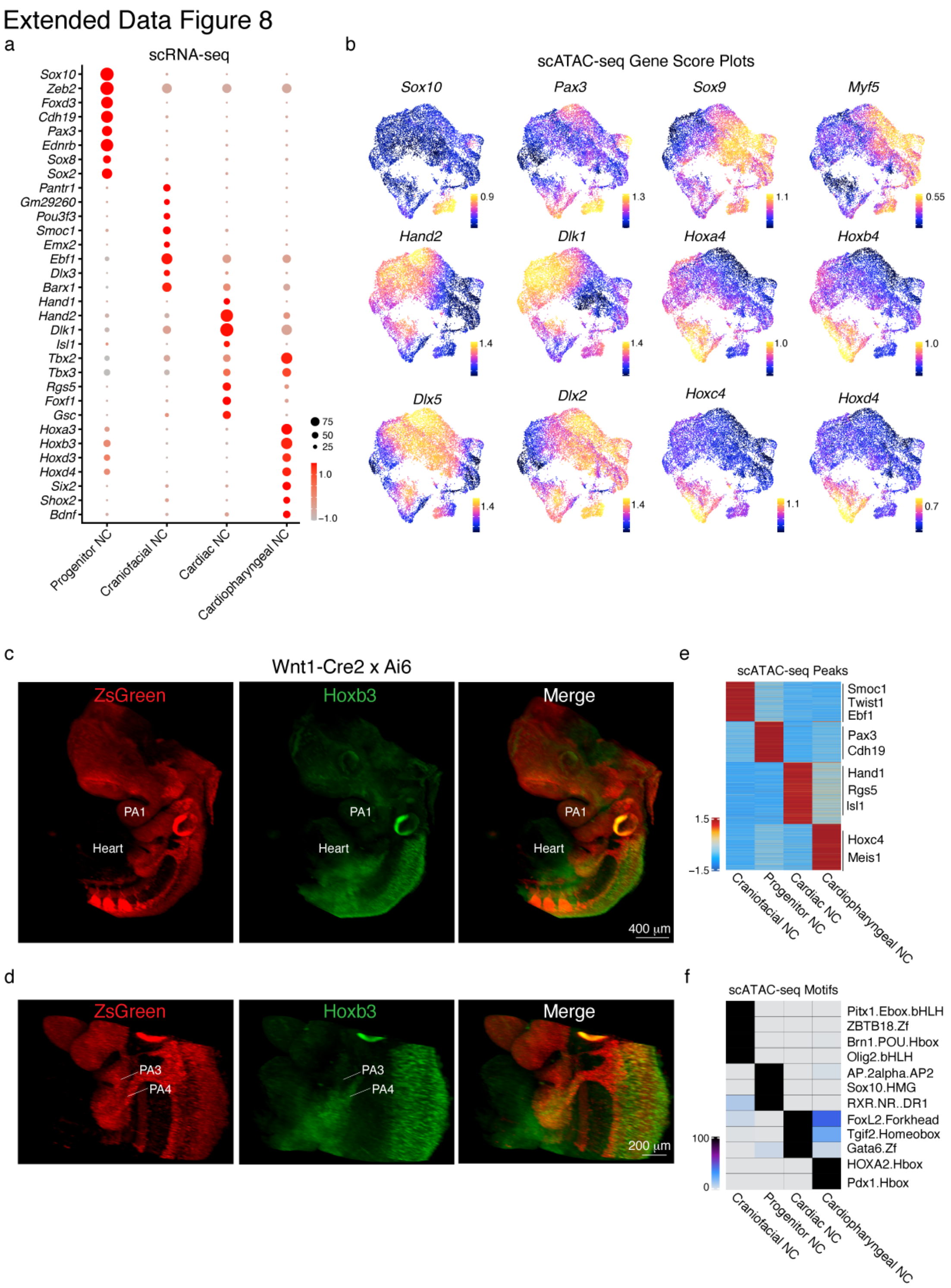


**Extended Data Fig. 8│ Identification of Four Broad Populations of Neural Crest Cells in Cardiopharyngeal Region at E9.5 – E11.5. a,** Dot plot of marker genes in four neural crest subpopulations. Color intensity represents scaled expression levels and dot size is proportion of cells within cluster expressing the gene. **b,** UMAP images of gene score values for representative marker genes within four neural crest subtypes (scATAC-seq), imputed values of chromatin accessibility within gene bodies and surrounding locus. Colors represent log2 normalized values + 1. **c,** Representative RNAscope images of wholemount E11.5 embryos from Wnt1-Cre2 crossed with Ai6 reporter mice and stained for ZsGreen and Hoxb3 (n=2). **d,** Zoomed-in view of same embryo, with focus on pharyngeal arches 3 and 4, where Hoxb3 expression overlaps with neural crest lineage. **e,** Heatmap of cluster-enriched peaks (FDR <= 0.5 & Log2FC >= 1.5). Colors represent z-scored values for enrichment of peak to a cluster. Genes listed on y-axis left were identified through HOMER. **f,** Heatmap of motifs enriched within each cluster using ArchR's peakAnnoEnrichment function with the HOMER database (FDR <= 0.5 & Log2FC >= 1). Color of cells scaled by normalized enrichment (-log10(p-adj)) ranging from 0-100.

Extended Data Figure 9

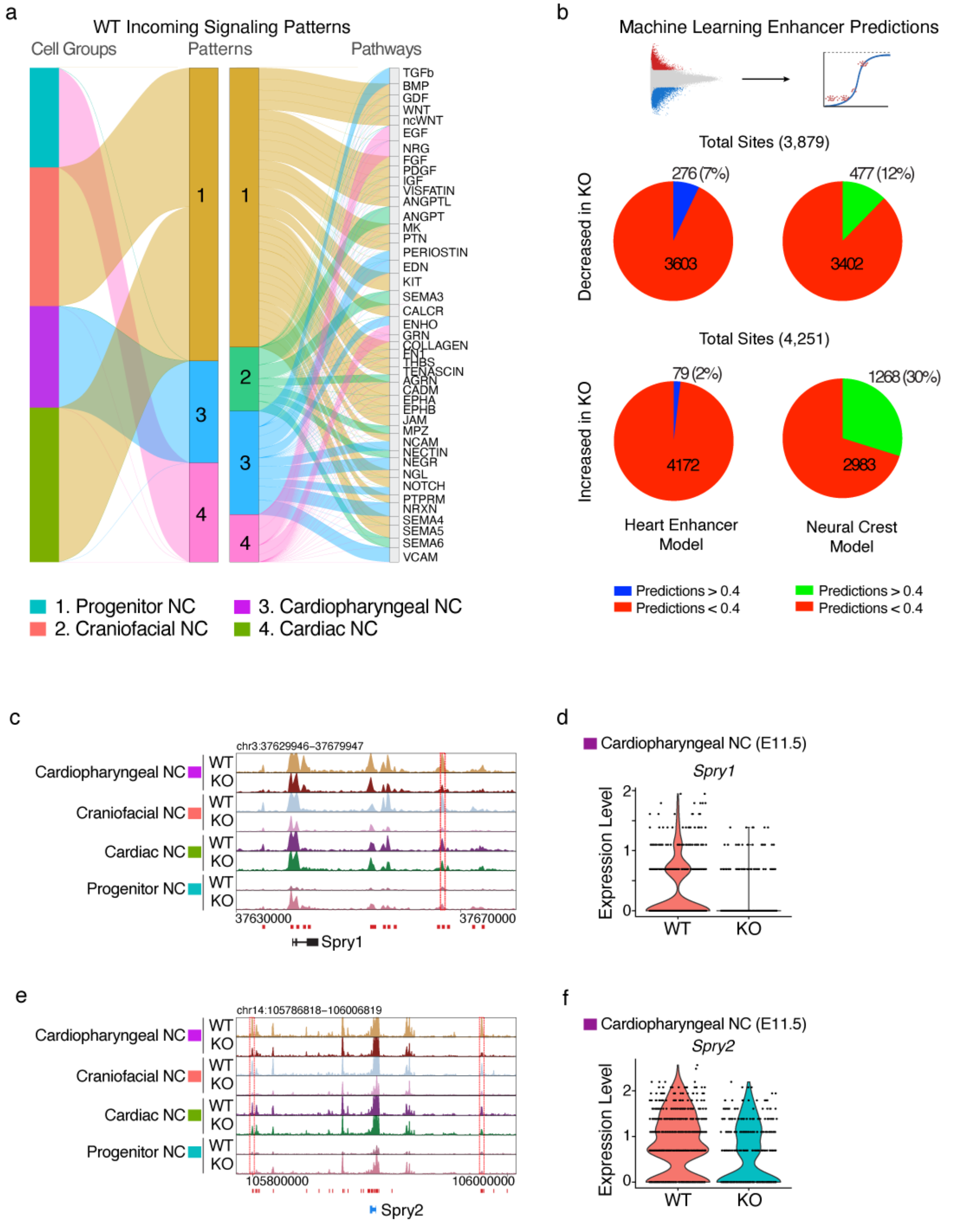


**Extended Data Fig. 9 | Differential Chromatin Accessibility in Non-Cardiac Neural Crest Populations. a,** River plot visualization of incoming signaling communication patterns (k=4 patterns) for WT neural crest cell types using Cellchat. Amongst the four patterns, three patterns were shown to be relevant and the signaling pathways within these patterns are shown on the right. **b,** Pie chart of intersection for filtering differentially accessible regions of cardiopharyngeal neural crest cells with both the cardiac and neural crest machine learning (prediction score cutoff > 0.4). **c,** Browser track view of *Spry1* gene locus and neural crest populations with coordinates for machine learning predicted enhancer at top and predicted peak highlighted in dotted red box. **d,** Violin plot of *Spry1* expression comparing WT and KO cells with cardiopharyngeal NCs at E11.5. **e,** Browser track view of *Spry2* gene locus and neural crest populations with coordinates for machine learning predicted enhancer at top and predicted peak highlighted in dotted red boxes. **f,** Violin plot of *Spry2* expression comparing WT and KO cells with cardiopharyngeal NCs at E11.5.

**Table S1. Metrics for scATAC-seq and scRNA-seq Samples**

**Table S2. Peak-to-Gene Linkage Coordinates**

**Table S3. Metrics for Zebrafish Enhancer Injections**
